# Microbial cross-feeding is stabilized when a dependent mutant is segregated from its independent ancestor

**DOI:** 10.1101/2025.01.23.634601

**Authors:** Olivia F. Schakel, Ryan K. Fritts, Anthony J. Zmuda, Sima Setayeshgar, James B. McKinlay

## Abstract

Microbial gene loss is hypothesized to be beneficial when gene function is costly, and the gene product can be replaced via cross-feeding from a neighbor. However, cross-fed metabolites are often only available at low concentrations, limiting the growth rates of gene-loss mutants that are dependent on those metabolites. Here we define conditions that support a loss of function mutant in a three-member bacterial community of: (i) N_2_-utilizing *Rhodopseudomonas palustris* as an NH^4+^ -excreting producer, (ii) N_2_-utilizing *Vibrio natriegens* as the ancestor, and (iii) a *V. natriegens* N_2_-utilizaton mutant that is dependent on the producer for NH^4+^. Using experimental and simulated cocultures, we found that the ancestor outcompeted the mutant due to low NH^4+^ availability under uniform conditions where both *V. natriegens* strains have equal access to nutrients. However, spatial structuring that separated the mutant from the ancestor, while maintaining access to NH^4+^ from the producer, allowed the mutant to avoid extinction. Counter to predictions, mutant enrichment under spatially structured conditions did not require a growth rate advantage from gene loss and the mutant coexisted with its ancestor. Thus, cross-feeding can originate from loss-of-function mutations that are otherwise detrimental, provided that the mutant can segregate from a competitive ancestor.

**Graphical abstract:** 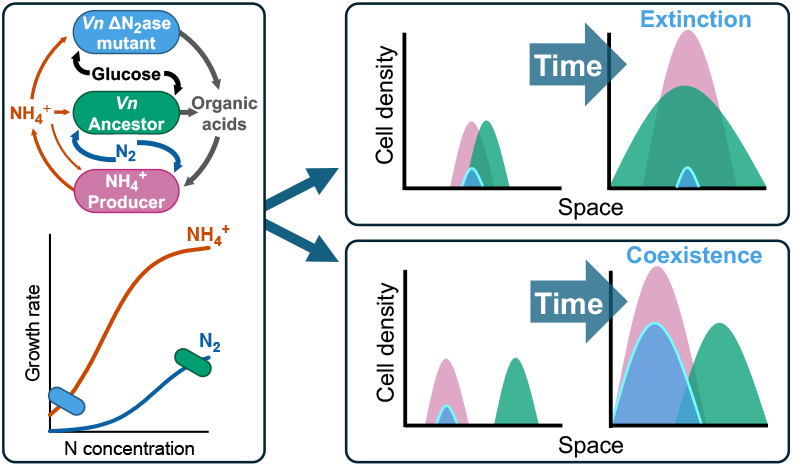

## Introduction

Individuals within microbial communities constantly adapt to changing environments.

One adaptation is beneficial loss-of-function (LOF) mutations, which are enriched when the cost of losing a gene is outweighed by the benefit of acquiring the gene product from a neighbor. This type of gene loss is perhaps best known as the Black Queen Hypothesis (BQH) [1, 2]. The BQH includes a ‘producer’ that creates a public good that promotes beneficial gene loss in a recipient ‘beneficiary’ [1, 2]. The term distinguishes beneficiaries from mutants with neutral or detrimental LOF mutations. Beneficiaries also do not harm the producer, distinguishing them from LOF ‘cheaters’ that exploit public goods at the expense of the producer [1, 3, 4]. The fitness advantage from the LOF mutation should lead to the extinction of the ancestral strain, provided that the producer is another species [1]. Adaptive gene loss supported by cross-feeding has been used to explain the natural prevalence of LOF mutants, like auxotrophs [5-9]. However, while nutrient-rich conditions are known to enrich for auxotrophs [10-13] and cross-feeding of molecules like iron-scavenging siderophores can enrich for cheaters [14-16], there are few direct observations where long-term cross-feeding led to the enrichment of spontaneously-evolved LOF beneficiaries [17, 18]; most studies used engineered LOF mutants.

One reason why gene loss might be infrequently observed is because cross-fed nutrients often exist at sub-saturating concentrations, preventing the mutant from achieving a maximum growth rate theoretically afforded by gene loss (Fig. 1). For example, a trait that was predicted to be subject to beneficial gene loss is N_2_ fixation, the conversion of nitrogen gas (N_2_) into ammonium (NH^4+^) via the cytoplasmic enzyme nitrogenase [1, 2]. N_2_ fixation is essential, expensive (e.g., 16 ATP per N_2_ fixed [19]), and NH^4+^ can passively escape the cell due to its equilibrium with membrane-permeable NH_3_ [19-22]. However, when we cocultured NH^4+^ - requiring *Escherichia coli* with wild-type (WT) N_2_-fixing *Rhodopseudomonas palustris, E. coli* initially grew at < 1% of the maximum possible rate and only reached 5% after 146 generations [22]. When we engineered *R. palustris* to excrete NH^4+^, *E. coli* still only grew at 25% or 43% of the maximum growth rate, depending on the level of NH^4+^ excretion [23]. Based on these observations, we predict that in the presence of an NH^4+^ -excreting producer, another N_2_-fixing bacterium would have a competitive advantage over a daughter nitrogenase LOF mutant, whose growth rate would be restricted by low NH_4_^+^ availability (Fig 1B, C).

**Figure 1.**
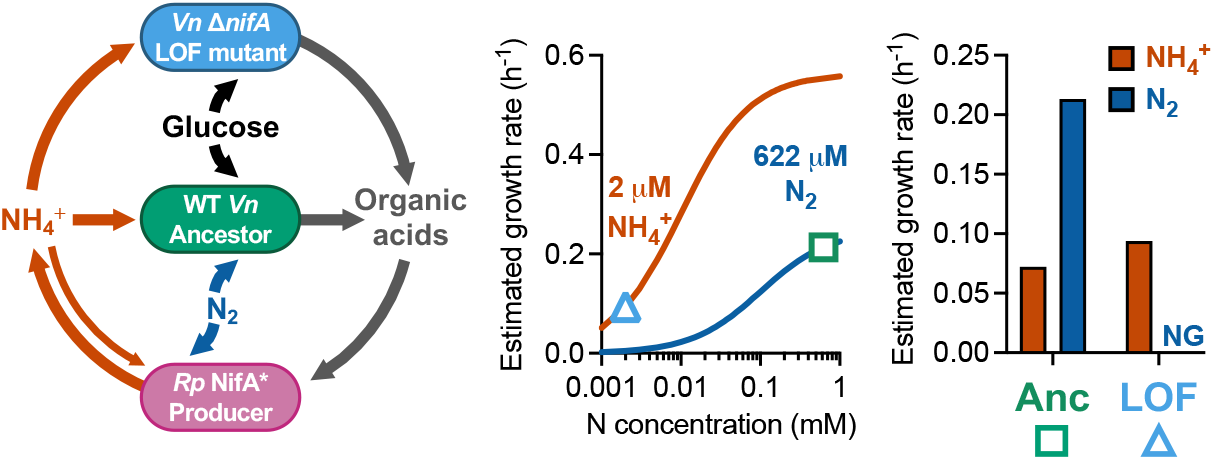
The LOF mutant growth rate is inferior to that of the ancestor due to the low concentration of cross-fed nutrient; (A) the coculture consists of (i) a producer, *R. palustris* (*Rp*) that fixes N_2_ and excretes NH_4_^+^ due to a NifA* mutation, and two *V. natriegens* (*Vn*) strains, (ii) a recipient that is incapable of N_2_ fixation and depended on the producer for NH_4_^+^, and (iii) a self-sufficient, N_2_-fixing ancestor; all strains are non-motile; (B) Monod model estimates of WT *V. natriegens* growth rates in the coculture based on the concentration of each nitrogen source (symbols); the growth rate with N_2_ (blue) would be higher than that with NH_4_^+^ (orange); see the Methods for model details; (C) estimated ancestor and recipient growth rates in coculture with each nitrogen source.

The emergence and maintenance of cross-feeding is also observed in some spatially structured populations [5, 24-30]. Community structure can create nutrient pockets, deserts, and gradients where populations have differential access based on local conditions [30-33].

Clustering of cooperating partners can decrease local nutrient concentrations, keeping cheaters to the fringes [24, 27, 29, 34-36]. Metabolite-externalizing populations can also achieve larger populations, despite carrying costly mutations, when physically aggregated within cross-feeding communities [28, 29]. Such spatially structured populations can accommodate diversity [37], including slow-growing subpopulations [33], cheaters [24, 29], and other competitors [38]. Thus, the extent to which fitness advantages from LOF mutations contribute to competitive outcomes in structured environments remains unclear.

Here we address conditions that can support the emergence of a nitrogenase LOF mutant. We used a defined bacterial community resembling one that could result in a BQH scenario to test whether the LOF mutant would emerge as a beneficiary. The community consisted of two species, where one was a NH_4_^+^ -excreting producer, and the other species was subdivided into a self-sufficient ancestor and a LOF mutant that was dependent on the producer (Fig. 1). Using both experimental cocultures and computational models, we found that the LOF mutant was always outcompeted by the ancestor under uniform conditions. However, we identified spatial conditions that allowed the LOF mutant to coexist with the ancestor, independently of any advantage afforded by the LOF mutation. Our results thus indicate that spatial structuring of populations can sustain LOF mutants without meeting BQH criteria of ancestor extinction nor a LOF mutation imparting a fitness advantage.

## Results

### Development of a coculture to test BQH predictions

Previously, we established obligate reciprocal cross-feeding between *E. coli* and an *R. palustris nifA*\* mutant [39]; *E. coli* fermented glucose and excreted organic acids as essential carbon for *R. palustris* and *R. palustris* fixed N_2_ and excreted NH_4_^+^ as essential nitrogen for *E. coli* [23]. N_2_ fixation was predicted to be subject to beneficial gene loss according to the BQH [1, 2]. To test whether loss of N_2_ fixation would be beneficial to bacteria cocultured with NH_4_^+^ -excreting *R. palustris*, we sought to replace *E. coli* with N_2_-fixing, fermentative, *Vibrio natriegens*. In the desired coculture, *V. natriegens* would comprise a self-sufficient ‘ancestor’ subpopulation and a recipient LOF mutant subpopulation that is dependent on the *R. palustris* ‘producer’ for NH_4_^+^ (Fig. 1A). We refrain from calling the LOF mutant a beneficiary unless we confirm a fitness benefit from the LOF mutation.

To prevent *V. natriegens* N_2_ fixation, we deleted *nifA*, encoding the transcriptional activator of nitrogenase genes. The Δ*nifA* LOF mutant did not grow with N_2_ but showed similar growth kinetics to the parent (ancestor) in monocultures and in *V. natriegens* cocultures with NH_4_Cl (Fig. S1; *R. palustris* omitted). We then verified that the producer could support the LOF mutant in coculture; population trends resembled *R. palustris* + *E. coli* cocultures, with both strains having a common exponential phase where the *V. natriegens* LOF growth rate more closely resembled that of *R. palustris nifA*\* than a monoculture growth rate [23] (Fig. S2A,B versus Fig. S1B,C). In contrast, coculturing *R. palustris* with the *V. natriegens* ancestor resembled *R. palustris* + *E. coli* cocultures with NH_4_Cl; rapid growth by the *V. natriegens* ancestor was followed by *R. palustris* growth [23] (Fig. S2C). Glucose was exhausted and organic acids accumulated in the first phase, indicative of *V. natriegens* ancestor growth, and then organic acids were depleted in the second phase, indicative of *R. palustris* growth (Fig. S2D). Having established the expected trends, we then examined cocultures comprised of the producer, ancestor, and LOF mutant (Fig. 1A).

### The LOF mutant is not enriched when *V. natriegens* subpopulations have equal access to nutrients

Like cocultures pairing the producer and ancestor (Fig. S2C), shaken cocultures combining the producer, ancestor, and LOF mutant had two growth phases (Fig. 2A). Tracking (sub)populations by colony-forming units (CFUs) showed that, aside from an early dominance by the LOF mutant, the ancestor outcompeted the LOF mutant by 24 h (Fig. 2B). At that time, glucose was exhausted (Fig. 2C), preventing further ancestor and LOF mutant growth.

**Figure 2.**
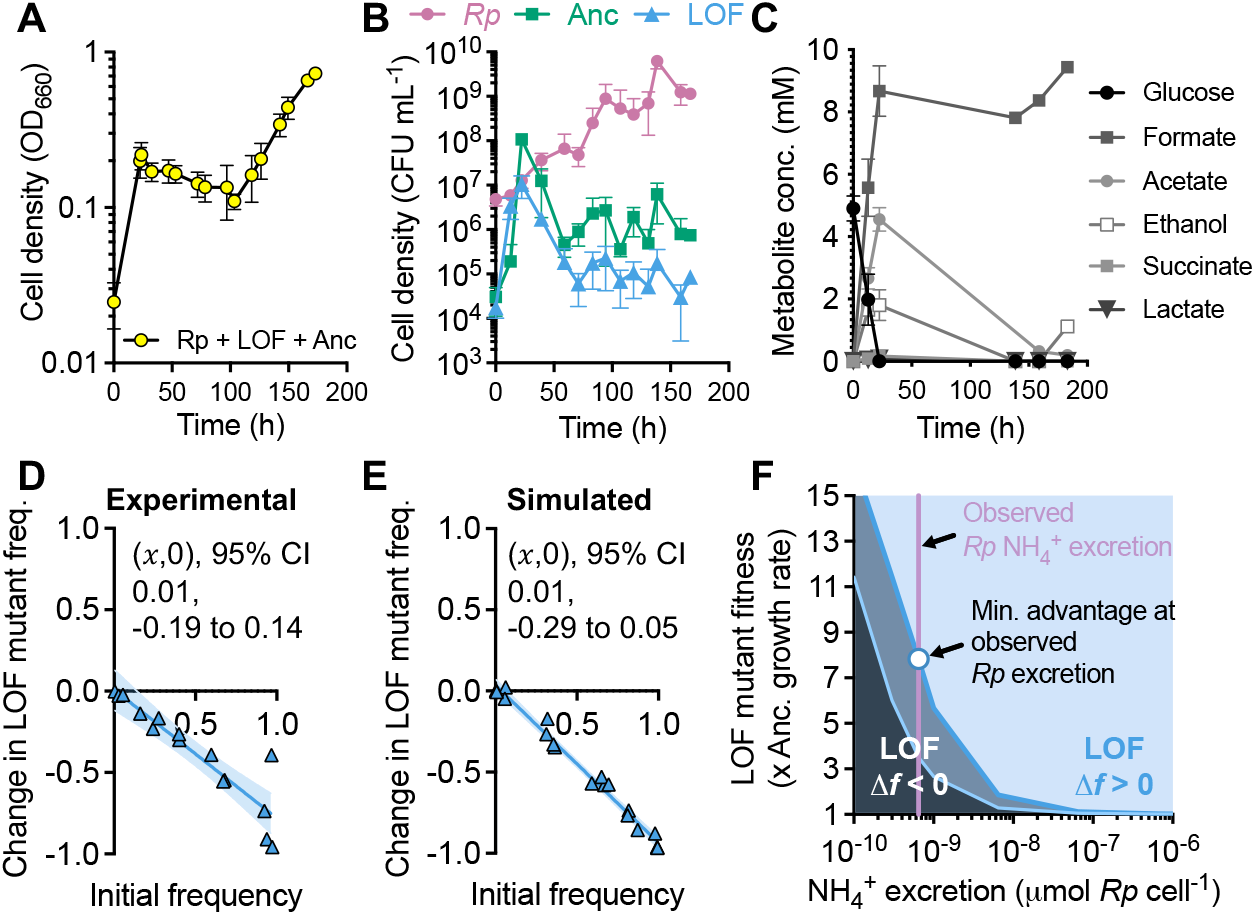
The ancestor outcompetes LOF mutant in shaken cocultures; (A) growth of shaken cocultures of *R. palustris, V. natriegens* Δ*nifA* LOF mutant, and WT *V. natriegens* ‘ancestor;’ (B) growth of each strain, determined using selective plating for CFUs; (C) glucose and fermentation product concentrations in cocultures; (A-C) points are means ± SD (n = 3); (D, E) IFR assays in shaken liquid experimental cocultures (D) or simulated cocultures with spatially uniform conditions (E); coexistence is assumed if the *x*-intercept (*x*,0) is between 0 and 1, otherwise it is assumed that one population drives the other to extinction; each point is a single experimental or simulated coculture; initial LOF mutant frequencies were the same for experimental and simulated cocultures (initial frequency range = 0.5% - 97.3% using *V. natriegens* populations only); *x*-intercept and 95% CI error bands were determined using linear regression analysis; (F) boundaries where the LOF mutant change in frequency goes from negative (Δ*f* < 0; extinction) to positive (Δ*f* > 0; enriched) for different producer (*Rp*) NH_4_^+^ excretion levels and LOF mutant maximum growth rate 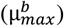 values relative to the ancestor (Anc); LOF mutant initial frequency *f*0 = 0.061; dark blue line, results with observed producer maximum growth rate; light blue line, results if producer maximum growth rate = ancestor maximum growth rate; purple vertical line, experimentally-estimated producer NH_4_^+^ excretion level; circle, minimum LOF mutant growth advantage required to avoid extinction at the experimentally-estimated producer NH_4_^+^ excretion level.

The above experiment used an initial LOF mutant frequency of ∼0.5, relative to the ancestor. To determine if the LOF mutant could be enriched from different initial frequencies we performed a reciprocal invasion-from-rare (IFR) assay, which also tests the mutual invasion criterion for coexistence [40, 41]. An *x*-intercept (*x*,0) between 0 and 1 suggests coexistence of the mutant and ancestor, whereas (*x*,0) ≥ 1 suggests ancestor extinction, and (*x*,0) ≤ 0 suggests mutant extinction [40-42]. IFR assays provide similar insights as serial transfers but they can be performed more quickly and are less prone to evolution affecting the results [41]. We inoculated the LOF mutant and ancestor at variable frequencies in shaken cocultures keeping the total initial *V. natriegens* population at a 1:1 ratio with *R. palustris*. The LOF mutant was consistently outcompeted by the ancestor ((*x*,0) = 0.01, 95% CI: -0.19 to 0.14; Fig. 2D).

We considered that LOF mutant enrichment could be influenced by NH_4_^+^ excretion level, the LOF mutant fitness advantage, and spatial heterogeneity. To explore these parameters, we built upon a mathematical model [23] to describe population growth and diffusion of cells and nutrients over a 2 × 2 cm domain (Supplementary materials, Table S1). We first tested whether the model could replicate experimental trends by simulating IFR conditions with uniformly distributed populations and nutrients. We gave the LOF mutant a maximum growth rate 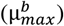 advantage of 1.1-times that of the ancestor, based on a comparison of *R. palustris* growth rates with and without nitrogenase expression (Fig. S3). The results resembled those from experimental cocultures with an *x*-intercept that was not significantly different from zero (Fig. 2E).

We then mapped what maximum growth rate 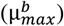 advantage would be required for the LOF mutant to have a positive change in frequency (Δ*f*) at different producer NH_4_^+^ excretion levels. At the experimentally-estimated NH_4_^+^ excretion level of 6.50 × 10^−10^ µmol cell^-1^ [23], the LOF mutant would need an unrealistic 8-fold maximum growth rate advantage over the ancestor (Fig. 2F, circle). The LOF mutant could be enriched with a lower growth rate advantage at NH_4_^+^ excretion levels 1-2 orders of magnitude higher than *R. palustris nifA*\* (Fig. 2F). While a wide range of NH_4_^+^ excretion levels can be engineered [23, 43, 44], natural examples are within an order of magnitude of *R. palustris nifA*\* excretion level (∼2 × 10^−10^ µmol cell^-1^ for *Azotobacter* [45] assuming a cell weight of 1 pg [46]). We also considered a scenario where the producer grows as fast as the ancestor (instead of a ∼3-fold difference in growth rate), to yield more public NH_4_^+^. Using the higher producer growth rate shifted the Δ*f* boundary in favor of LOF mutant enrichment, but the required growth advantage was still ∼3.5-times that of the ancestor at the experimentally-estimated NH_4_^+^ excretion level (Fig. 2F).

We also considered how NH_4_^+^ privatization might affect population outcomes. NH_4_^+^ from nitrogenase is highly privatized; N_2_ fixation occurs in the cytoplasm where most NH_4_^+^ will be assimilated before it can escape. Generation of public goods outside the cell can profoundly affect producer-consumer relationships [2, 47-50]. Low privatization can enrich for LOF cheaters [14, 34, 51-53] and thus might also enrich for LOF beneficiaries. To explicitly address low privatization, we modified our model to describe a hypothetical production of NH_4_^+^ outside of the cell by both N_2_ fixing bacteria [47] (Fig. 3A). In this scenario, the LOF mutant was predicted to outcompete the ancestor, provided the LOF mutant had a growth rate advantage (Fig. 3B). Thus, while our results suggest that N_2_ fixation is unlikely to lead to the enrichment of LOF beneficiaries, the outcome could be different for less-privatized public goods. Moving forward, we focused on whether spatial structuring of populations might facilitate LOF nitrogenase mutant enrichment, despite high privatization.

**Figure 3.**
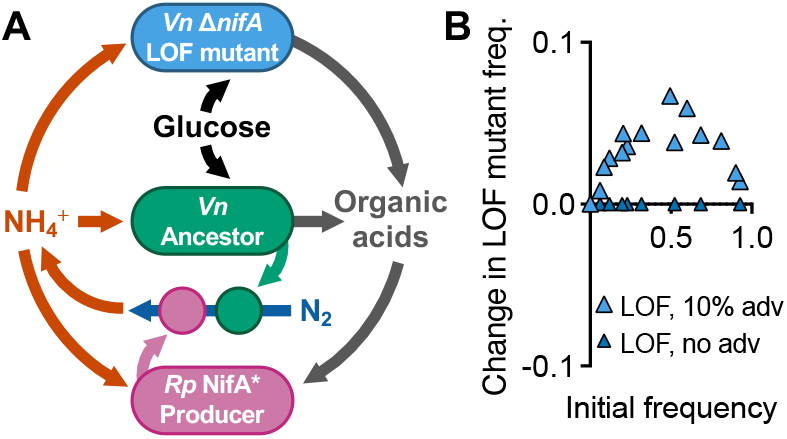
Hypothetical extracellular NH_4_^+^ production allows for enrichment of the LOF mutant in accordance with the BQH; (A) the modified model allows for production of NH_4_^+^ via a hypothetical extracellular enzyme (circles) produced by the N_2_-fixing producer and ancestor populations; (B) simulated IFR with extracellular NH_4_^+^ production under spatially uniform conditions; initial LOF mutant frequencies were the same for experimental and simulated cocultures (range = 0.1% - 92.8% using *V. natriegens* populations only).

### Static conditions do not enrich for the LOF mutant

Spatially structured communities can foster populations that might otherwise have a fitness disadvantage [28, 29, 33, 38]. We thus tested whether spatial structuring could lead to nitrogenase LOF mutant enrichment. We began with minimal intervention by coculturing non-motile strains under static conditions in either liquid or a fluid 0.15% agarose matrix (Fig. 4A); we assumed that random spatial structuring developed as the cells settled. Our IFR assays, sampled after mixing at the final time point, suggested that the LOF mutant would go extinct; both conditions gave *x*-intercepts that were not significantly different from zero (Fig 4B). We also tested these conditions using the mathematical model with a random initial cell distribution and diffusion constants consistent with agarose, where appropriate (Table S1). The simulations also suggested LOF mutant extinction in both conditions (Fig. 4C).

**Figure 4.**
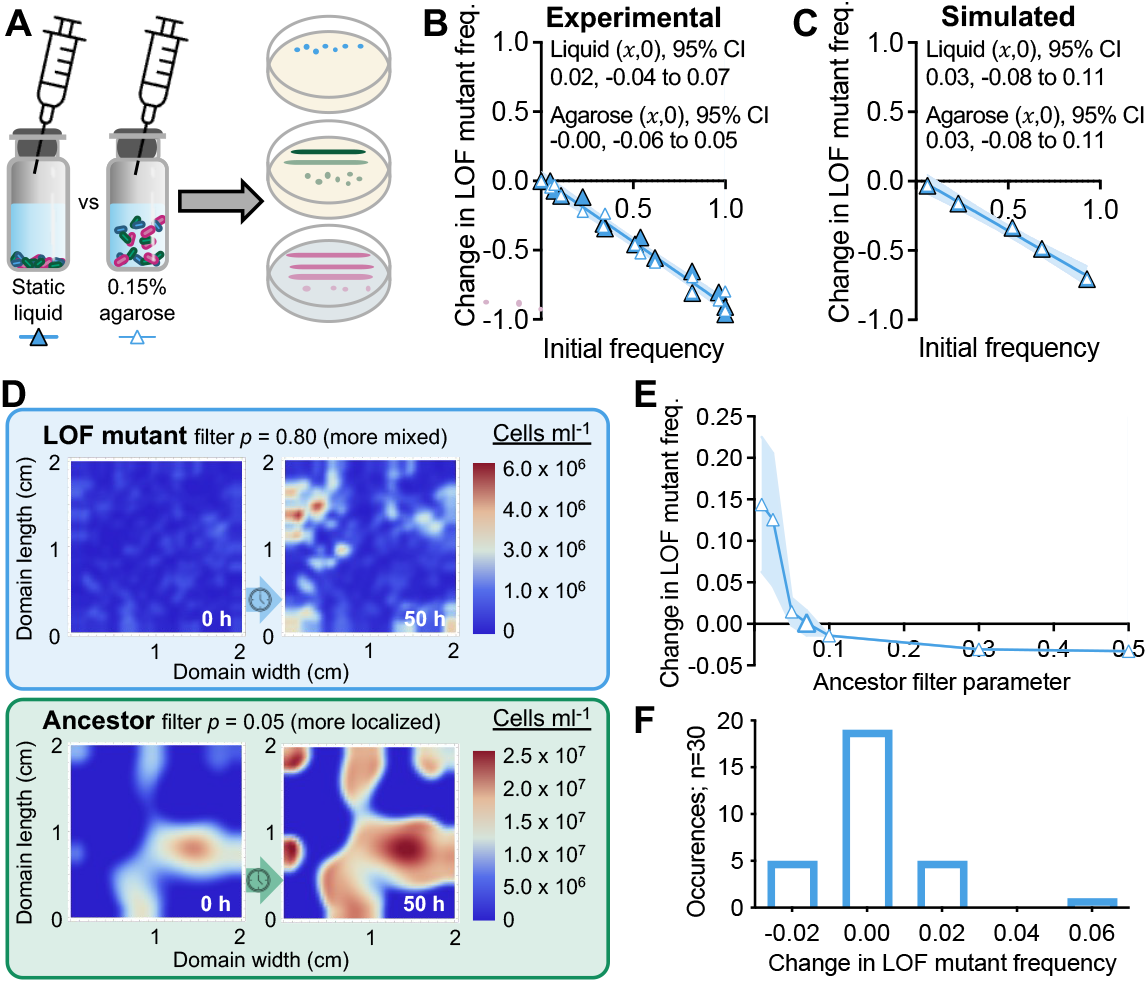
Isolation of ancestor populations can theoretically lead to local and domain-level enrichment of the LOF mutant; (A) experimental approaches to randomly distribute non-motile cells in static cocultures without or with a fluid 0.15% agarose matrix; community composition was assessed after 6 d by selective plating; experimental (B) and simulated (C) IFR assays with randomized cell distributions in static liquid or 0.15% agarose; each point is from an individual experimental or simulated coculture; initial LOF mutant frequencies were the same for experimental and simulated cocultures (initial frequency range = 0.1% - 99.9% using *V. natriegens* populations only); experimental IFR assays were only mixed before sampling; *x*-intercept and 95% CI error bands were determined using linear regression analysis; simulated agarose used lower diffusion coefficients; (D) simulated cell densities in a 2 × 2 cm domain (assumed to be uniform across height) at 0 h and 50 h; initial LOF mutant frequency *f*0 was 0.061; initial beneficiary and producer populations were colocalized using a common filter parameter (*p* = 0.80); the ancestor population was distributed using *p* = 0.05; (E) mean LOF mutant change in frequency (Δ*f*(*t*)) from different random cell distributions (LOF mutant and producer, *p* = 0.80 for all ancestor filter parameter values); n = 10 except the enlarged data point, where n = 30; error bands = SD; (F) histogram for the enlarged data point in (E) where the ancestor *p* = 0.07 (n=30), and the LOF mutant change in frequency is ∼ 0; this threshold change in frequency value is unique to these simulation parameters.

### Segregation leads to LOF mutant enrichment without a maximum growth rate advantage

Cross-feeding neighborhoods can occur at the scale of one to several cells [31]. Thus, while we did not observe LOF mutant enrichment at the domain level, there could have been pockets of local enrichment. We addressed this possibility using the model by varying a spatial filter (*p*) for the initial ancestor random distribution, while maintaining a spatial filter value for the initial random LOF mutant and producer distributions (*p* = 0.8). Larger filter parameters give rise to a more fine-grained spatial variation, resulting in more mixing of populations (Fig. S4). Smaller filter parameters correspond to spatial coarsening, creating regions where the ancestor is more isolated (e.g., *p* = 0.05, Fig. 4D). Simulations using *p* = 0.05 showed that the LOF-mutant can expand its population in regions where the ancestor population remained low (Fig. 4D). When averaged across the entire domain, the LOF mutant could be enriched when the initial ancestor distribution was coarse (*p* ≤ 0.07; Fig. 4E, F).

To directly address how segregation impacts population outcomes, we simulated Gaussian distributions of each initial population at distinct sites (Fig. 5A; ancestor at (*x* =0.5, *y* =1 cm) and the LOF mutant and producer colocalized at (*x*=1.5, *y*=1 cm). Upon glucose depletion (∼150 h), the ancestor had grown around the LOF mutant and producer populations, highlighting that the populations did not grow in compartmentalized isolation (Fig. 5A). Despite some spatial overlap of the initial Gaussian distributions, segregation led to LOF mutant enrichment ((*x*,0) = 0.38; 95% CI: 0.36 to 0.43), with predicted coexistence with the ancestor (Fig. 5B).

**Figure 5.**
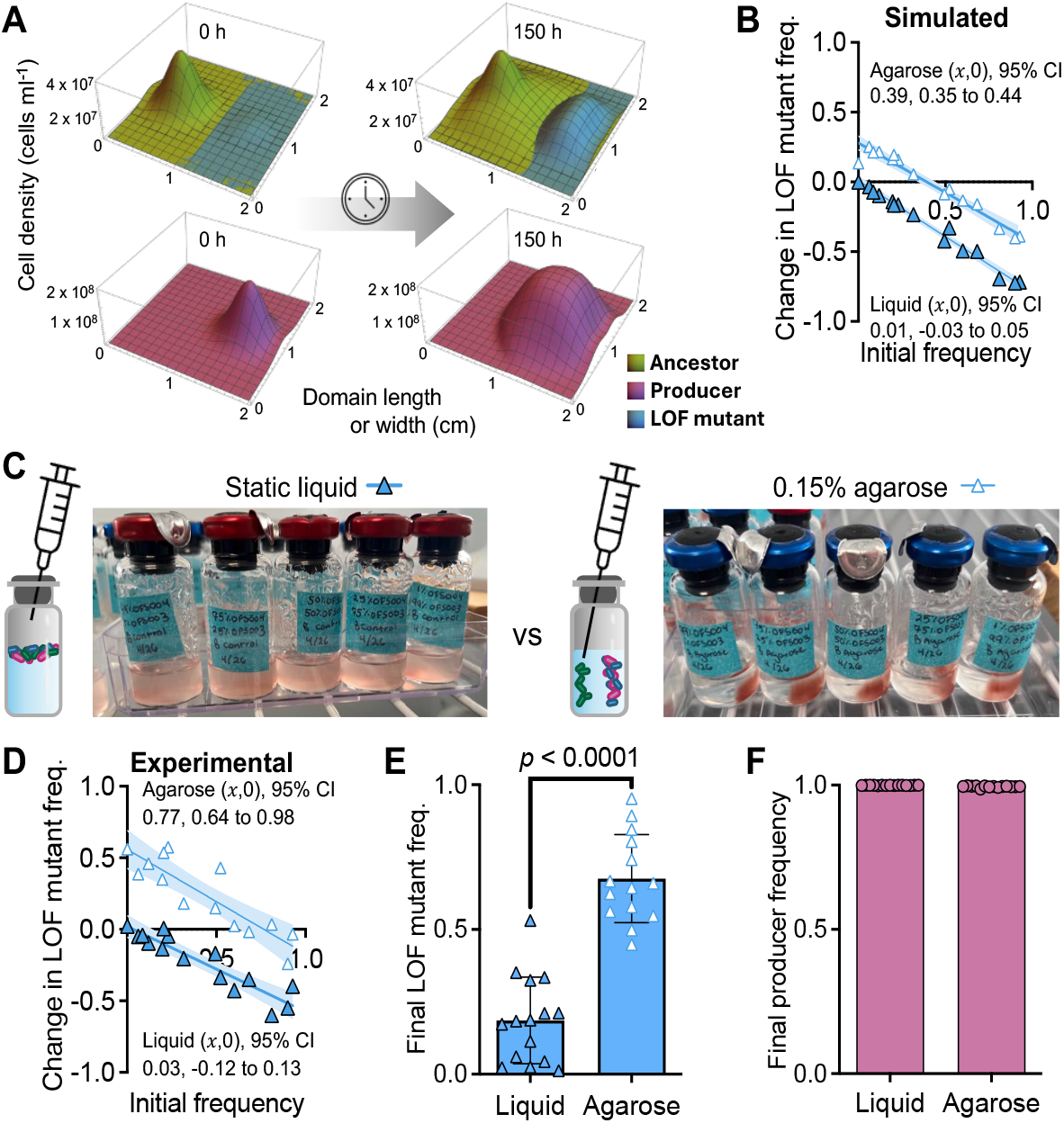
Colocalization with the producer and separation from the ancestor leads to domain-level enrichment of the LOF mutant; (A) example of simulated Gaussian distributions of initial populations; the LOF mutant and the producer were colocalized but are shown on separate graphs for visualization purposes; cell densities are assumed to be uniform across height; (B) simulated IFR in liquid versus 0.15% agarose using the distinct inoculation sites in (A); (C) experimental ccocultures were inoculated to static liquid or to locations in 0.15% agarose; localization is evident from pigmented *R. palustris* growth after 6 days; (D) IFR from the experimental conditions; cocultures were only mixed before sampling (B, D) each point is from an individual experimental or simulated coculture; initial LOF mutant frequencies were the same for experimental and simulated cocultures (initial frequency range = 0.1% - 92.7% using *V. natriegens* populations only); *x*-intercept and 95% CI error bands were determined using linear regression analysis; final LOF mutant frequency (E, using *V. natriegens* populations only) and the producer (F, using all populations) in static liquid or 0.15% agarose; bars are means ± SD, n=15; *p*-value is from a two-tail t-test.

Motivated by these simulations, we constructed an experimental counterpart by inoculating populations at specific locations just below the surface of 0.15% agarose media (Fig. 5C). We first verified that populations were localized, but not fully isolated, using non-motile ancestor monocultures; the ancestor was detected at the opposite side of the vial by 20 h, spreading via diffusion and growth (Fig. S5). Glucose was also depleted more slowly on the opposite side of the vial (Fig. S5). Thus, inoculating populations on opposite sides of the vial should allow for interactions, but with less local competition.

We then co-inoculated the producer and the LOF mutant on one side of the vial and the ancestor on the other side (Fig. 5C). In agreement with the simulations, the LOF mutant was enriched, with predicted coexistence with the ancestor ((*x*,0) = 0.77; 95% CI: 0.64 to 0.98; Fig. 5D). The final LOF mutant frequency was also significant compared to in static liquid (Fig. 5E, *V. natriegens* populations only; the producer was the dominant species, making up ∼99% of the total population, Fig. 5F).

Coexistence differs from the BQH prediction of ancestor extinction [1]. We thus investigated whether our results met the BQH criterion of a LOF mutant fitness advantage [1], for which we used growth rate. Although the simulated NH_4_^+^ levels exceeded the half-maximum value (*km*) by an order of magnitude (Fig. S6), the highest growth rate 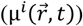 at any location in the spatial domain for the LOF mutant never exceeded that of the ancestor (Fig. 6A, S6).

**Figure 6.**
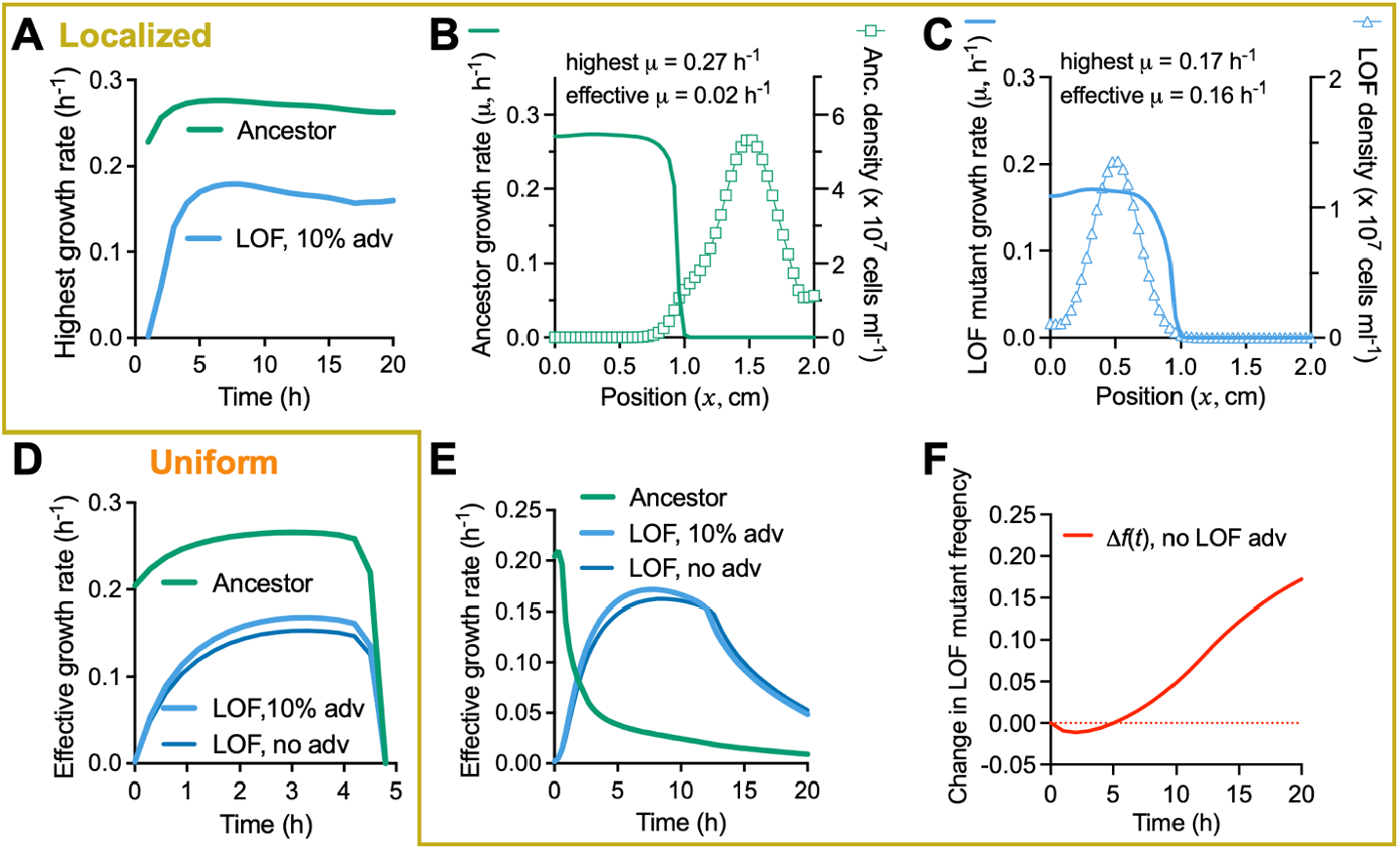
Separation from the ancestor allows the LOF mutant to be enriched without an intrinsic maximum growth rate 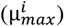 advantage when colocalized with the producer; all graphs are from simulated cocultures using an initial LOF mutant frequency of 6.1% (*V. natriegens* populations only); all localized conditions (A-C,E,F) used initial population spatial distributions as in Fig 5 B, (σ = 0.2) (A) the highest growth rates in the spatial domain; (B, C) a cross-section across the domain at t = 10 h and *y* = 1 cm shows that the highest growth rate does not always occur at the same location as the highest cell density; to compare growth rates, we therefore adopted an effective growth rate 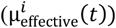, which is a spatially averaged growth rate weighed by cell density (Supplementary material); effective growth rates with uniform (D) of localized conditions (E) when the LOF mutant does, and does not have a 10% maximum growth rate advantage (adv) over the ancestor (Anc); (F) change in frequency for the conditions in (E) when the LOF mutant does not have a maximum growth advantage; change in frequency was calculated from the integrals of the effective growth rates (Supplementary material. Fig. S7).

Using growth rate as the fitness metric in numerical simulations with non-uniform spatial conditions can be misleading; the calculated growth rate at a given point in time and space can be high because of nutrient availability but cells might be absent to take advantage of local conditions (Fig. 6B, C). We therefore examined effective growth rate 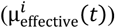, a spatially averaged growth rate weighted by cell density (Fig. 6, Supplementary materials). Under uniform conditions, the effective ancestor growth rate was always higher than that of the LOF mutant (Fig. 6D). However, for localized conditions, the LOF mutant achieved a higher effective growth rate for most of the simulation, explaining its enrichment (Fig. 6E). To assess whether the encoded LOF mutant maximum growth rate 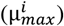 advantage, afforded by the LOF mutation, contributed to this outcome, we simulated the same conditions with no maximum growth advantage. In this case, the effective LOF mutant growth rate was lower, but not enough to affect population outcomes (Fig. 6D-F; change in fitness was determined from the time integral of the effective growth rate; Fig. S7). Thus, our results suggest that the LOF mutant enrichment was due to the initial segregation from the ancestor, leading to local glucose depletion that prevented invasion by the ancestor (Fig. S6). In other words, spatial conditions, rather than an intrinsic fitness advantage from gene loss, led to the enrichment of the LOF mutant.

### LOF mutant enrichment does not require producer colocalization

LOF mutant enrichment could have been due to (i) separation from the ancestor and/or (ii) producer colocalization. To assess the contribution of each factor, we simulated inoculation sites as above and increased population overlap by modifying the standard deviation of the Gaussian distributions (σ) for all populations (Fig. 7A). Increasing σ led to a decrease in the IFR *x*-intercept (Fig. 7B); for σ > 0.4, IFR plots resembled uniform conditions, highlighting the importance of LOF mutant separation from the ancestor.

**Figure 7.**
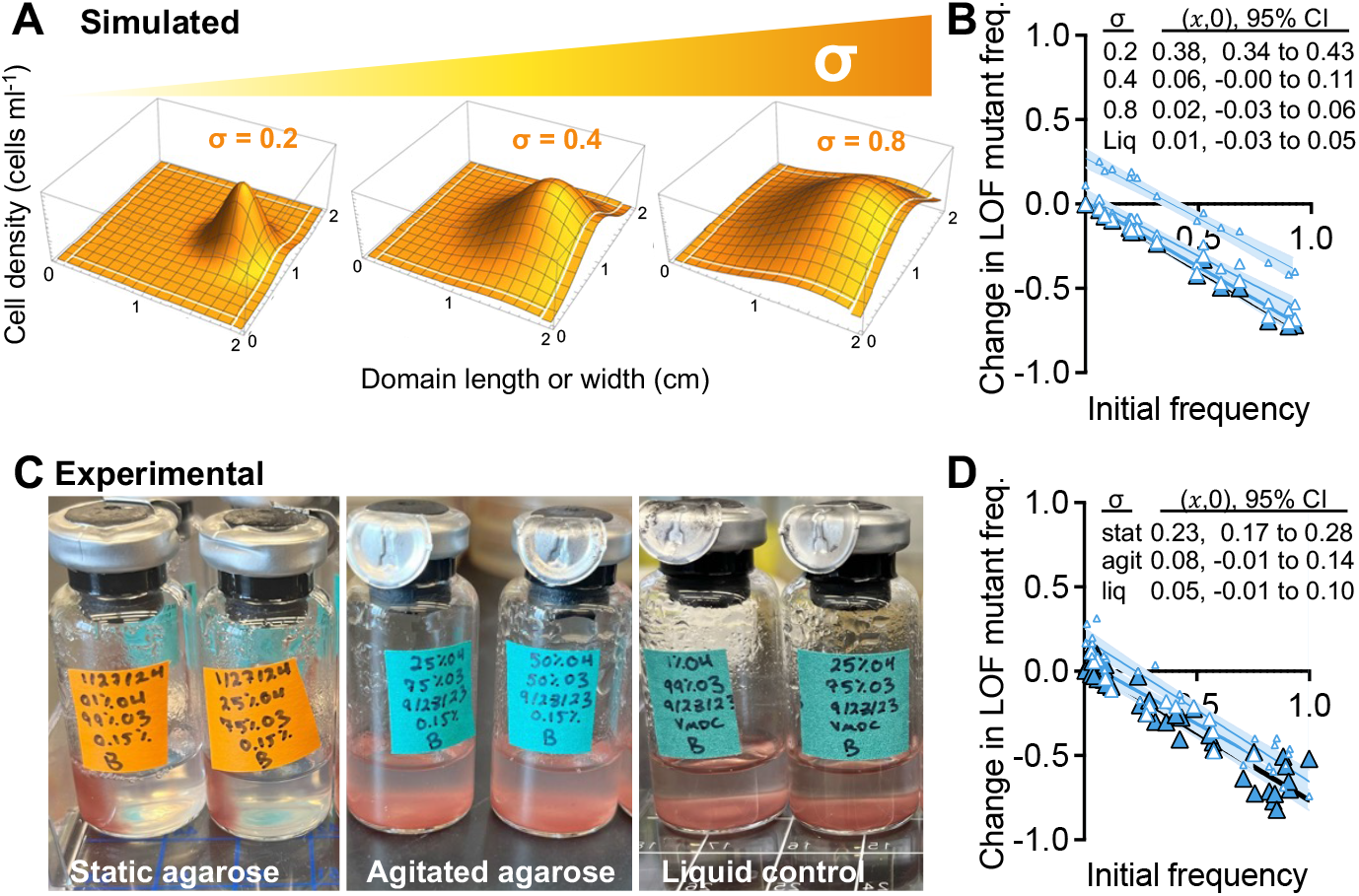
Increasing population overlap prevents enrichment of the LOF mutant; (A) initial population distribution varied by σ (standard deviation of Gaussian distributed inoculum); (B) simulated IFR assays using different σ for all populations to affect overlap between the ancestor and LOF populations; inoculum locations were the same as in Fig. 5, where the LOF mutant and producer are colocalized; (C) experimental conditions varied σ by not disturbing 0.15% agarose (stat), agitating agarose by stirring before inoculation (agit), or using static liquid (liq); the degree of localized populations is evident from growth of pigmented *R. palustris* after 6 days; (D) IFR results from (C) cocultures were only mixed before sampling; (B, D) each point is from an individual experimental or simulated coculture; initial LOF mutant frequencies were the same for experimental and simulated cocultures (initial frequency range = 3.1% - 91.4% using *V. natriegens* populations only); *x*-intercept and 95% CI error bands were determined using linear regression analysis.

To experimentally test these simulated predictions, we agitated 0.15% agarose by stirring to fragment the polymer and thereby widening initial distributions at inoculation sites (Fig. 7C). Consistent with the simulated results, agitating the agarose (increasing σ), moved the IFR results to resemble uniform conditions (Fig. 7D). The sensitivity of the matrix to disturbance might also explain the different values from different IFR assays (Fig 5D vs Fig 7D) compared to static liquid that always gave *x*-intercept values that were not significantly different from zero (Fig. S8). Our results indicate that less spatial overlap with the ancestor is essential for LOF mutant enrichment, even when the mutant and producer are colocalized.

To assess the role of LOF mutant and producer colocalization in LOF mutant enrichment, we simulated IFR conditions with the producer and ancestor colocalized and separated from the LOF mutant (Fig. 8A; σ = 0.2). The *x*-intercept again suggested LOF mutant coexistence with the ancestor, but at a lower equilibrium frequency ((*x*,0) = 0.06; 95% CI: 0.03 to 0.10; Fig 8B). This result was confirmed experimentally by inoculating populations in a similar spatial arrangement in 0.15% agarose ((*x*,0) = 0.07; 95% CI: 0.04 to 0.19; Fig. 8C). Again, this result was not dependent the LOF mutant maximum growth rate advantage (Fig. 8D). Overall, our results suggest that separation from the ancestor is more important for LOF mutant enrichment than producer colocalization, provided there is still sufficient access to NH_4_^+^ from the producer.

**Figure 8.**
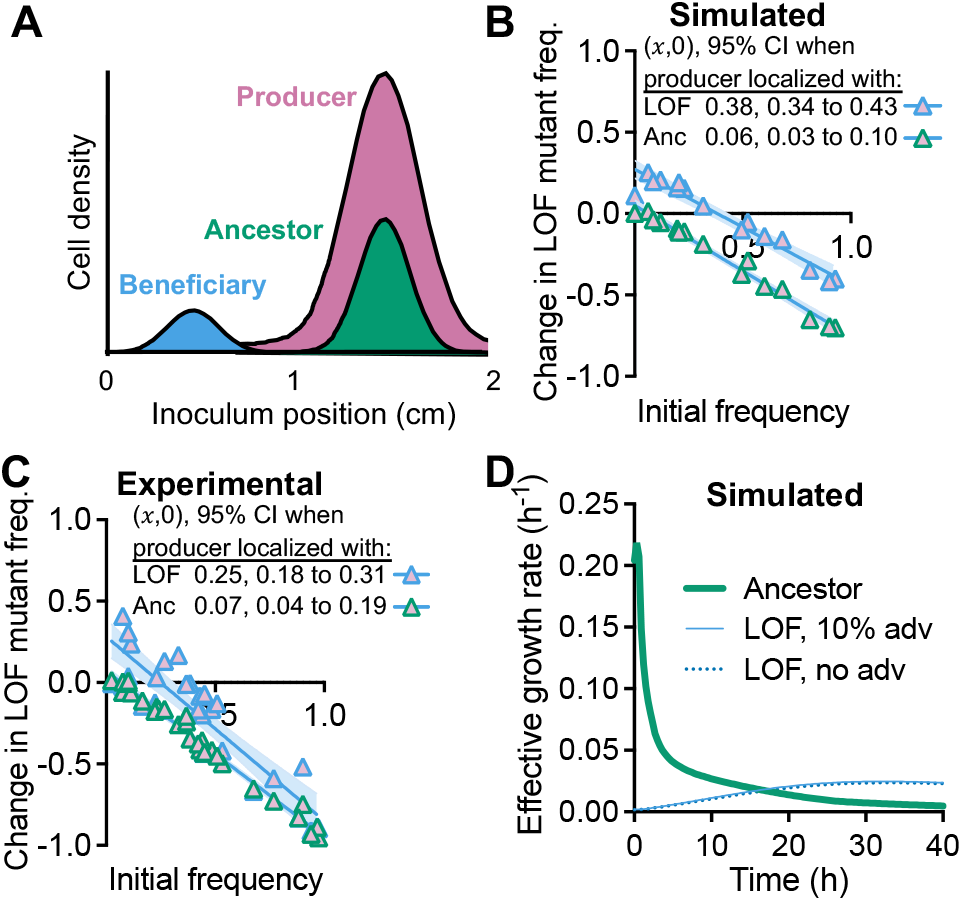
The LOF mutant can be enriched even when distant from colocalized ancestor and producer populations; (A) initial population distributions (not to scale); (B) simulated and experimental; (C) IFR assays comparing conditions where either the LOF mutant or the ancestor is colocalized with the producer; each point is from an individual simulated or experimental coculture; initial LOF mutant frequencies were the same for experimental and simulated cocultures (initial frequency range = 0.1% - 92.8% using *V. natriegens* populations only); experimental cocultures were only mixed before sampling; *x*-intercept and 95% CI error bands were determined using linear regression analysis; (D) effective growth rates for a simulation where the initial populations are distributed as in (A) with an initial LOF mutant frequency *f*0 = 0.061.

## Discussion

We used an experimental coculture and mathematical model to test the BQH prediction that loss of nitrogenase would be beneficial [1]. Beneficial nitrogenase loss did not seem feasible given that low extracellular NH_4_^+^ would prevent a higher growth rate than that with N_2_ (Fig. 1). Indeed, the LOF mutant was consistently outcompeted by the ancestor under uniform conditions (Fig. 2, Fig. S6). However, spatial separation from the ancestor, while maintaining access to NH_4_^+^ from the producer, led to LOF mutant enrichment (Fig. 4-8). Thus, although we observed nitrogenase mutant enrichment, the outcome differed from BQH predictions in two ways [1]: (i) the data suggested mutant coexistence with the ancestor (Fig. 4-8; IFR (*x*,0) between 0 -1), and (ii) a maximum growth rate 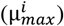 advantage from gene loss was not required (Fig. 6, 8).

### Partial privatization and LOF mutant outcomes

There are likely other cases where sub-saturating cross-fed nutrients under uniform conditions would favor the ancestor over a LOF mutant. However, the privatization level of a cross-fed resource could flip this outcome, even under uniform conditions [2, 47-50, 52, 53]. For example, Lerch et al predicted that intracellularly-generated NH_4_^+^ from N_2_ fixation is less likely to support LOF mutants than extracellular iron-scavenging siderophores, for which ancestors and LOF mutants have equal access [48]. This finding agrees with our hypothetical scenario where we simulated extracellular NH_4_^+^ production and observed LOF mutant dominance (Fig. 3). Privatization might help explain why there are few reports of spontaneous auxotroph emergence during long-term cross-feeding of intracellularly generated compounds; in one example, emergent amino acid auxotrophs appeared to be transient [10]. However, in examples of less-privatized detoxification services, spontaneous LOF mutants appeared to be stable [17, 18]. Still, engineered pairings of amino acid auxotrophs and producers suggest that beneficial fitness outcomes are possible [54-56]. One possibility is that these cases involved easily overlooked but important spatial organization, like microscopic cell clusters [28].

### Microbial community structure can accommodate disadvantageous gene loss

The nitrogenase LOF mutation in our study is clearly disadvantageous. Thus, the mutant in this case is neither a cheater nor a beneficiary. LOF mutant enrichment instead depended on separation from its competitive ancestor. Our findings resemble others where spatial structuring prevented direct competition with LOF cheaters [24, 27, 29, 34-36], but with an important difference. Whereas a LOF cheater would emerge within a large ancestor population that is exploitable, our LOF nitrogenase mutant would emerge within a large ancestor population that is competitively superior. How can a LOF mutant with a disadvantage escape its ancestor?

Physical separation can be achieved through dispersal of the LOF mutant or the ancestor. For example, three bacterial species cocultured in a nutrient rich environment could only coexist in a biofilm if there was biased dispersal of the dominant competitor [57]. Whereas we used non-motile strains to provide more control over location, microbial motility can also influence cooperator-competitor interactions. For example, competition against a cheater was improved when an amino acid cross-feeding bacterium was motile [29]. Motility also allowed for slower-growing cooperative strains to increase in frequency relative to a cheater in swim agar [58]. Non-motile cells can also escape communities via fluid flow (advection) [59]. Transient flow across a surface can rapidly isolate cells, albeit at a low frequency in lab conditions [60]. Other forms of community disruption or bottlenecking can also lead to physical separation and benefit a slow-growing subpopulation [61, 62].

With prolonged separation from the ancestor and access to a cross-feeding partner, obligate dependencies through additional LOF mutations could evolve [5, 6], and perhaps contribute to genome streamlining, which the BQH can partially address [1, 63, 64]. But what mechanisms would enrich for additional LOF mutations? Although separation from competitors can lead to the enrichment of a LOF mutant, it seems improbable that each mutation would coincide with separation events that also maintain access to a cross-feeding partner. More likely, separation from the ancestor is just one factor contributing to the origin and maintenance gene loss-associated cross-feeding, along with mutations that impart competitive advantages, like those described by the BQH. Our findings underscore that environmental features that influence spatial community structure are important to consider in the evolution of cooperative phenotypes that might otherwise seem to defy evolutionary theory.

## Methods

### Bacterial strains

Strains, plasmids, and primers are in Table S2-4. Mutations were verified via Sanger sequencing. *V. natriegens* strains were derived from TND1964, containing pMMB-tfoX [65] (WT). *V. natriegens* mutations were made by introducing linear PCR products via natural transformation [65, 66]. The ‘ancestor,’ OFS003, is a kanamycin resistant (Δ*dns*::Kan^R^), non-motile (Δ*flgE*; flagellar hook) derivative of the WT strain (Fig. S9). The LOF mutant, OFS004, additionally carries a Δ*nifA* mutation, preventing N_2_ fixation, and *dns* is instead replaced by a spectinomycin resistance cassette. Each cassette had a comparable effect on the growth rate (Fig. S10).

The ‘producer,’ *R. palustris* CGA4067, is a non-motile derivative of CGA4005 [47], which itself is derived from type strain CGA0092 [67]. CGA4067 is incapable of H_2_ oxidation (Δ*hupS*), has low cell aggregation [68], excretes NH_4_^+^ due to a mutation in *nifA* [39], and is non-motile due to deletion of *motAB* (flagella stator). CGA4039 was made incapable of N_2_ fixation by deleting structural genes for all three nitrogenase isozymes [69]. *R. palustris* deletion mutations were made by homologous recombination after introducing the appropriate suicide vector [69, 70], via electroporation [71] as described [23, 72].

### Growth conditions

Cultures were incubated at 30°C with light from a halogen bulb (750 lumens). Anoxic media was prepared by bubbling media with N_2_ in culture vessels, then sealing with rubber stoppers and aluminum crimps prior to autoclaving.

*V. natriegens* and *R. palustris* were recovered from 25% glycerol frozen stocks (−80°C) on agar plates containing LB3 (lysogeny broth with 2% w/w NaCl) for *V. natriegens* or photosynthetic media (PM) [73] with 10 mM disodium succinate for *R. palustris*. Kanamycin (100 ug/mL) or spectinomycin (200 ug/mL) were included where appropriate. Starter cultures were grown from single colonies. Starter cultures were grown in 27-ml anaerobic tubes with 10 ml of minimal media: *R. palustris* was grown in M9-derived coculture media (MDC) [23] with 1.5 mM disodium succinate. *V. natriegens* was grown in MDC modified with (final concentrations): 10 mM glucose, 80 mM NaCl, 200 mM MOPS (pH 7), and 0.5 mM NH_4_Cl, to gently transition cells to N_2_-fixing conditions; this media, without NH_4_Cl, is called VMDC. A 1% inoculum of *R. palustris* and a 0.5% inoculum of each *V. natriegens* strain was then used to start anaerobic cocultures in VMDC with 5 mM glucose.

Shaken (150 RPM) cocultures, including for IFR assays, were grown in 10-ml volumes in 27-ml tubes, oriented horizontally. Static IFR assays were performed in 4-ml volumes in 10-ml anaerobic serum vials with or without agarose (Research Products International). Contaminating nitrogen was removed from agarose (Fig. S11) by washing 0.15g of agarose twice with ultra-pure water and then once with VMDC (12-ml volumes in a 15-ml conical tube; agarose was pelleted by centrifuging at 2415 x g and removing supernatants by pipette). Washed agarose was resuspended in 100 ml VMDC in a 160-ml serum vial before making anoxic and autoclaving.

After autoclaving, molten agarose was kept suspended during cooling by rocking overnight (Boekel Scientific). Agitated agarose (Fig. 7) was stirred with a stir bar overnight during cooling (200 RPM). Glucose and cations were added and then agarose media was then dispensed into serum vials by syringe using a 1”, 23-gauge needle (BD). IFR assays were inoculated with 9 × 10^6^ cells of producer and 9 × 10^6^ cells of LOF mutant plus ancestor at the specified ratio. For randomized static cell distributions, the inoculum was dropped onto the media and allowed to settle during incubation. Localized populations were inoculated on opposite sides of the vial using a 2”, 21-gauge needle (BD), to slowly inject cells just below the surface, without touching the glass.

### Analytical procedures

Motility was determined by using a pipette tip to stab a single colony into LB3 with 0.3% agar and then measuring swim diameter 17 h later. Cell density was measured as optical density at 660 nm (OD660) using a Genesys 20 spectrophotometer (Thermo-Fisher) or CFUs on selective media (see above). Growth rates were determined by fitting an exponential trendline using Microsoft Excel. Glucose, organic acids, and ethanol were quantified using a Shimadzu high performance liquid chromatograph as described [74]. For IFR assays, initial cell densities in agarose were assumed to be the same as those determined in 0.3 ml samples from liquid controls. Final cell densities, and metabolite levels were determined after 6 days by vortexing vials and then sampling 1 ml. For location sampling, 0.35 ml was taken using a 2”, 21-gauge needle (Fig. S5); cultures were then discarded. LOF mutant change in frequency Δ*f* = (LOF /(LOF + ancestor))final – (LOF /(LOF + ancestor))initial [41]. Linear regression for IFR assays and other statistical analyses were performed using Graphpad Prism v10.

### Mathematical modeling

The Monod model (Fig. 1) was: *μ* = *μ*MAX • S / [S + km]), where: *μ*, growth rate; *μ*MAX, maximum growth rate; S, NH_4_^+^ or N_2_ concentration; km, half-saturation constant for S. Parameter values: NH_4_^+^, 2 µM based on 20 µM / *R. palustris* cell [23]; N_2_, 622 µM based on Henry’s law assuming 1.02 atm and N_2_ solubility of 6.1*10^−4^ mol/(L*atm) [75]; *μ*MAX with NH_4_^+^, 0.43 h^-1^; *μ*MAX with N_2_, 0.25 h^-1^; kmNH_4_^+^, 0.01 mM [76]; kmN_2_, 0.1 mM [77]. A 1.3-fold *μ*MAX advantage with NH_4_^+^ was assumed for the LOF mutant based on a comparison of *R. palustris* strains (Fig. S3).

Population and metabolic dynamics in cocultures were simulated in Mathematica (v13.3 Wolfram Research, Inc., 2023) using coupled, nonlinear reaction-diffusion equations modified from previous models describing cross-feeding between *R. palustris* and *E. coli* [23, 47]. We numerically simulated the equations subject to no-flux boundary conditions using Mathematica’s NDSolve [r] function employing a stiff solver. Default parameter values are given in Table S1. Cell densities, *c*_*i*_, are in number of cells per ml, and the numerical solution corresponds to time-dependent concentrations in a system size of (*L*_*x*_ = 2 cm) × (*L*_*y*_ = 2 cm2 × (1 cm). The time evolution of cell densities in the *x* − *y* plane is investigated under different conditions, assuming the concentrations in the *z* −direction are uniform. Diffusion constants for cells and nutrients in liquid media versus the agarose matrix were estimated based on the Stokes-Einstein relation (Table S1). Model equations and details are available in the Supplementary material.

## Acknowledgements

We thank A. Stoner for assistance constructing CGA4039 and N. Haas for constructing NH003 and NH004. We thank C. Love for piloting early experiments and M. Di Salvo and K. Locey for useful discussions during the initial stages of the project. We also thank the McKinlay lab and the local Vibrio group for helpful advice.

## Data Availability

All data for this study are in this article and its supplementary information files.

