## Supplementary material for "Microbial cross-feeding is stabilized when a dependent mutant is segregated from its independent ancestor"

Short title: Rival segregation stabilizes cross-feeding

Olivia F. Schakel[1], Ryan K. Fritts[1a], Anthony J. Zmuda[1b],
Sima Setayeshgar[2]*, and James B. McKinlay[1]*
[1]Department of Biology, Indiana University, Bloomington, IN, USA
[2]Department of Physics, Indiana University, Bloomington, IN, USA

*Corresponding authors:
Setayeshgar: 727 E 3rd Street, Bloomington, IN 47405, USA;
McKinlay: 1001 E 3rd Street, Bloomington, IN 47405, USA;

Current address:
[a] Department of Molecular, Cellular and Developmental Biology and Cooperative Institute for Research in Environmental Sciences, University of Colorado Boulder, Boulder, CO, 80309, USA
[b] Department of Plant and Microbial Biology, University of Minnesota, Twin Cities, Saint Paul, MN, 55108, USA

**Mathematical model of cross-feeding.**
To numerically investigate the enrichment of a loss-of-function (LOF) mutant via mutualistic cross-feeding in bacterial cocultures, we constructed a parsimonious model of the cross-feeding between a slow-growing $N_2$-fixing cooperator (producer), paired with fast-growing fermentative partners that can either also fix $N_2$ (ancestor), or that lost $N_2$-fixation (LOF mutant) thereby requiring the product, ammonium ($NH_4^+$). The mathematical model, given by Eqs. 1-12, is a system of coupled, nonlinear reaction-diffusion equations, for three bacterial strains and nutrient levels given by organic acids, $N_2$, $NH_4^+$, and glucose, based on earlier work experimentally quantifying a mutualism between $N_2$-utilizing and $NH_4^+$-excreting *Rhodopseudomonas palustris* and $NH_4^+$-requiring *Escherichia coli* [1]. A minimal modification of this model to address non-privatized access to $NH_4^+$ production is given by Eqs. 14-18. Our experimental platform utilizes *R. palustris* as the $NH_4^+$-producer, and *Vibrio natriegens* as both the LOF mutant and the ancestor of the LOF mutant subpopulation, where the ancestor and can fix $N_2$ whereas the LOF mutant (*ΔnifA*) cannot fix $N_2$ and requires $NH_4^+$.

We numerically simulated the equations in two dimensions subject to no-flux boundary conditions using Mathematica's NDSolve [r] function employing a stiff solver (Wolfram Research, Inc., Mathematica, Version 13.3, Champaign, IL (2023)). Parameter values and diffusion constants are in Table S1. Rate constant dimensions are $h^{-1}$. Cell densities, $c_i$, are in number of cells per ml, and the numerical solution corresponds to time-dependent concentrations in a system size of $(L_x = 2\text{ cm}) \times (L_y = 2\text{ cm}) \times (1\text{ cm})$. The role of spatial localization of cell densities in the $x - y$ plane is investigated, assuming the concentrations in the $z$ −direction are uniform.

***Bacterial growth in all models is given by the following Monod functions*** [2]:

LOF mutant growth rate ($h^{-1}$) (1)

$$\mu^b(G, A) = \mu^b_{max} \frac{G}{G + K^b_G} \frac{A}{A + K^b_A},$$

Ancestor growth rate on $NH_4^+$ ($h^{-1}$) (2)

$$\mu^a_A(G, A) = \mu^a_{max,A} \frac{G}{G + K^a_G} \frac{A}{A + K^a_A},$$

Ancestor growth rate on $N_2$ ($h^{-1}$) (3)

$$\mu^a_N(G, N, A) = \mu^a_{max,N} \frac{G}{G + K^a_G} \frac{N}{N + K^a_A} \left(1 - \frac{A}{A + K^a_A}\right),$$

Producer growth rate on $NH_4^+$ ($h^{-1}$) (4)

$$\mu^p_A(C, A) = \mu^p_{max,A} \frac{C}{C + K^p_G} \frac{A}{A + K^p_A},$$

Producer growth rate on $N_2$ ($h^{-1}$) (5)

$$\mu^p_N(N, C, A) = \mu^p_{max,N} \frac{C}{C + K^p_G} \frac{N}{N + K^p_A} \left(1 - \frac{A}{A + K^p_A}\right).$$

Additionally, in the numerical implementation, the growth terms are multiplied by a Heaviside function of cell densities, imposing a cut-off of 1 cell $ml^{-1}$. Related to this cut-off, in the stochastic Fisher-Kolmogorov-Petrovsky equation, which represents a broad class of models that yield nonlinear traveling waves, it has been shown that fluctuations in the wave tip are important in determining the resulting wave speed [3, 4]. Extensions of our deterministic model aim to similarly explore stochastic effects, expected to be important in regions where cell numbers are low.

***The partially-privatized model uses the following equations to describe changes in population and metabolite levels:***

Organic acid concentration (mM): (6)

$$\frac{\partial C}{\partial t} = D_C \nabla^2 C + (F_C(\mu^b(G,A)c_b + \mu_A^a(G,A)c_a) + F_{C,N}\ \mu_N^a\ (G,N,A)c_a - (\mu_A^p(C,A)/Y_{C,A} + \mu_N^p(N,C,A)/Y_{C,N})c_p$$

Glucose concentration (mM): (7)

$$\frac{\partial G}{\partial t} = D_G \nabla^2 G - \mu_A^b(G,A)c_b/Y_G - (\mu_A^a(G,A)/Y_G + \ \mu_N^a(G,N,A)/Y_{G,N}^a)c_a$$

$N_2$ concentration (mM): (8)

$$\frac{\partial N}{\partial t} = D_G \nabla^2 N - \mu_N^b(G,N,A)c_a/Y_N^a - \frac{\mu_N^p(N,C,A)c_p}{Y_N^p}$$

$NH_4^+$ concentration (mM): (9)

$$\frac{\partial A}{\partial t} = D_A \nabla^2 A + F_A^p \mu_N^p(N,C,A)c_P + \ F_A^a \mu_A^a(G,N,A)c_a \ - c_p \mu_A^p(C,A)/Y_A^p - c_a \mu_A^a(G,A)/Y_A^a - c_b \mu^b(G,A)/Y_A^b$$

LOF mutant cell density (cells $ml^{-1}$) (10)

$$\frac{\partial c_b}{\partial t} = D_b \nabla^2 c_b + \mu^b(G,A)c_b$$

Ancestor cell density (cells $ml^{-1}$) (11)

$$\frac{\partial c_a}{\partial t} = D_a \nabla^2 c_a + (\mu_A^a(G,A) + \mu_N^a(G,N,A))c_b$$

Producer cell density (cells $ml^{-1}$) (12)

$$\frac{\partial c_p}{\partial t} = D_p \nabla^2 c_p + (\mu_A^p(C,A) + \mu_N^p(N,C,A))c_p$$

***The non-privatized model uses the following equations to describe changes in population and metabolite levels.***

We address non-privatized access to $NH_4^+$ produced by the producer and the ancestor, in this case achieved via a hypothetical extracellular $NH_4^+$-producing enzyme, as a minimal modification to the partially-privatized model equations. As part of minimizing the number of modifications, entities for $N_2$ and the extracellular $NH_4^+$-producing enzyme are not explicitly shown, and instead we assume that (i) $N_2$ is available at a constant saturating concentration, thus allowing Eq. 8 to be omitted, (ii) the concentration of $NH_4^+$-producing enzymes is proportional to populations of $N_2$-fixing organisms, and (iii) the rate of $NH_4^+$-production by the enzyme is constant and functions independently of population growth rates. Eqs. 6-12 are thus replaced with the following:

Organic acid concentration (mM): (13)

$$\frac{\partial C}{\partial t} = D_C \nabla^2 C + F_{C,A}(\mu^b(G,A)c_b + \mu_A^a(G,A)c_a) - \mu_A^p(C,A)/Y_{C,A}$$

Glucose concentration (mM): (14)

$$\frac{\partial G}{\partial t} = D_G \nabla^2 G - \mu_A^b(G,A)c_b/Y_G - \mu_A^a(G,A)/Y_G$$

$NH_4^+$ concentration (mM): (15)

$$\frac{\partial A}{\partial t} = D_A \nabla^2 A - c_p \mu_A^p(C,A)/Y_A^p - c_a \mu_A^a(G,A)/Y_A^a - c_b \mu^b(G,A)/Y_A^b + \widetilde{F_A^p} c_P + \widetilde{F_A^a} c_a$$

LOF mutant cell density (cells $ml^{-1}$) (16)

$$\frac{\partial c_b}{\partial t} = D_b \nabla^2 c_b + \mu^b(G,A)c_b$$

Ancestor cell density (cells $ml^{-1}$) (17)

$$\frac{\partial c_a}{\partial t} = D_a \nabla^2 C_a + \mu_A^a(G,A)c_a$$

Producer cell density (cells $ml^{-1}$) (18)

$$\frac{\partial c_p}{\partial t} = D_p \nabla^2 c_p + \mu_A^p(C,A)c_p$$

***Initial conditions with localized cell distributions.***

For spatially localized initial conditions, we use Gaussian distributions with given mean and standard deviation to specify the spatial localization of cell densities:

$$c_i(x,y) = c_{0_i} e^{-\frac{(x-x_i)^2 + (y-y_i)^2}{2\sigma_i^2}}. \quad (19)$$

The normalization $c_{0_i}$ is determined to achieve a specified total initial cell count $\mathcal{N}_i(0)$ for species $i$ for a given system size, where initial total cell numbers in simulations are chosen to be consistent with those in the experimental chamber. Populations can be weakly or sharply localized (large or small $\sigma_i$, respectively), and the extent of co-localization is additionally determined by the mean positions, $(x_i, y_i)$. To directly compare invasion from rare plots, where the initial frequency and the change in frequencies (horizontal and vertical axes) are specified in terms of the LOF mutant and ancestor cell numbers, the total producer population must be the same. Therefore, to compare with experimental invasion from rare measurements, in simulations we use initial producer cell numbers that are the same as those in experiments for each initial frequency.

***Initial conditions with random cell distributions.***
To generate random initial cell density distributions, we start with pink noise in two dimensions, which has a power spectrum of $1/k^{\alpha}$ (α = 2 in two dimensions), where $k = 2\pi/\lambda$ is the wavenumber, and λ is the wavelength. Pink noise is ubiquitous in nature and describes the statistical fluctuations of diverse physical and biological systems. The statistics of the random distribution used to generate the initial cell densities is not relevant for this analysis; rather, the focus is to address in a systematic way how successively coarsening the initial random distribution of the ancestral strain, thereby going from spatially "well-mixed" to "localized" cell densities, impacts emergence of the LOF mutant. Specifically, the pink noise random distribution is low pass filtered, attenuating wave numbers above a specified cut-off (or equivalently, attenuating the short wavelength components of the pink noise distribution). The numerical values are first clipped at zero (to achieve cell densities ≥ 0), smoothed with a Gaussian filter (to avoid sharp cell density gradients as initial conditions to the numerical simulations) and normalized to give desired total cell counts in the spatial domain. In this way, we systematically generated initial spatial density distributions displaying less point-to-point spatial variation by decreasing the filter or "fine-graining" parameter, $p$. The spatial coarsening that results from low pass filtering the ancestral distribution gives rise to larger regions of space where the LOF mutant is not proximal to the ancestor.

Fig. S4 shows representative initial cell densities for the LOF mutant and producer strains (Fig. S4A top row; filter parameter, $p = 0.8$) and initial cell density for the ancestor using different values of the filter parameter, demonstrating the transition from greater spatial variability (for larger filter parameter values) to lesser spatial variability (for smaller filter parameter values) (Fig. S4A, bottom row). When the LOF mutant, producer, and ancestor initial cell densities have greater spatial variability (e.g., $p = 0.8$) the populations are well-mixed and the change in LOF mutant frequency is negative (Fig. 4E, Fig. S4C), consistent with spatially uniform initial cell densities. When the ancestor is initially more localized, the LOF mutant is enriched (Fig. 4D, E; Fig. S4B).

***Diffusion in agarose media.***

Previous experimental work addressed diffusion of various solutes and bacterial cells in biofilms and porous media, relevant as the natural habitat for the cross-feeding dynamics considered here [5-9]. The Stokes-Einstein relation for the diffusion constant of a sphere of radius R in a low Reynolds number fluid, is given by:

$$\mathrm{D} = \frac{k_B T}{6\pi\eta\mathrm{R}}\,, \qquad (20)$$

where $k_B$ is the Boltzmann constant, $T$ is temperature and $\eta$ is the fluid viscosity. While this description well represents diffusion of solutes and nonmotile cells in bulk fluid, the diffusion constant of motile cells is larger and not captured by this result [5, 6]. As a benchmark, for a sphere of radius $R \sim 1$ μm, $T = 25^{o}$C and $\eta = 0.001$ Pa · sec (water), $D \sim$ 2:2 x $10^{-9}$ $cm^2$ $s^{-1}$. For rod shaped cells modeled as an ellipsoid, $R \longrightarrow R/log(2R/b)$, where $R$ and $b$ are the semimajor and semiminor axes, respectively [7]. For a rod-shaped cell that is 2 μm in length and 1 μm in diameter, this introduces a factor of ~ 1.4 increase in $D$. At $30^{o}$C, the viscosity of water is given by $\eta = 0.0008$ Pa · sec, so for diffusion of nonmotile cells in bulk fluid at the coculture temperature, we find $D \sim 3.8$ x $10^{-9}$ $cm^2$ $s^{-1}$. In our simulations, for all strains, we use $D_{cell} \sim 5$ x $10^{-9}$ $cm^2$ $s^{-1}$ = 1.8 x $10^{-5}$ $cm^2$ $h^{-1}$. For the diffusion constant of small metabolites in water, ranging from $10^{-10}$ to $10^{-9}$ $cm^2$ $s^{-1}$, we use 8 x$10^{-10}$ $cm^2$ $s^{-1}$ ~ 0.03 $cm^2$ $h^{-1}$.

Within a biofilm matrix, which represents a tortuous medium, diffusion is slower. Previous work measured the reduced effective diffusion constant of selected microbial solutes in biofilms [8], demonstrating a decrease by a factor of 0.2 to 0.8. In experimental studies of the rheological properties of agarose fluid gels, it has been shown that in the low agarose concentration regime, the increase in viscosity with agarose concentration is approximately linear [9]. For a 0.5% agarose solution, the viscosity at $30^{o}$C can be extracted from these measurements to be approximately 0.004 Pa · sec, which is ~5 times greater than the viscosity of water, leading to diffusion constants that correspondingly smaller by this factor [9]. These experimental results are consistent with measurements of diffusion constants in biofilms. For our numerical simulations in agarose, accounting for the increase in viscosity, we use diffusion constants that are 3-times smaller than in bulk fluid (Table S1).

***Connection with Fisher-KPP equation.***

The Fisher-KPP equation, due separately to Fisher [10] and Kolmogorov, Petrovsky, and Piscounov [11] is a nonlinear reaction-diffusion equation which in one dimension (Fisher's equation) can be written as

$$c_t = Dc_{xx} + \alpha c\left(1 - \frac{c}{K}\right). \tag{21}$$

The Fisher-KPP equation supports travelling wave solutions used to model spread of various invasive phenomena [12-15]. From phase plane analysis of the traveling wave solution, $c(x,t) = c(z), z = x - vt,$ it can be shown that for any $v > 0$, there exists a unique right traveling wave connecting the state $(c = K, c_x = 0)$ for $x \rightarrow -\infty$ to the state $(c = 0, c_x = 0)$ for $x \rightarrow \infty$. For $v \geq v_c$, where the critical wave speed is $v_c = 2\sqrt{\alpha D}$, the wave is nonoscillatory and a monotonically decreasing function of $x$. From Figure S5, which demonstrates spread of the ancestor in monoculture inoculated at one side of the chamber to the opposite end (in 0.15% agarose), we approximate a wave speed of $v \sim 2$ cm per 150 h or 0.013 cm $h^{-1}$. For a monotonically decreasing travelling front, requiring $v \geq v_c$ leads to $\alpha \leq v/4D$. Using our values for diffusion of cells in agarose, $D{\sim}6\ x10^{-6}$ $cm^2$ $h^{-1}$, and $v \sim 0.013$ cm $h^{-1}$, we find that with a growth rate for the ancestor (*V. natriegens* grown anaerobically in minimal media with glucose) given by $\alpha \leq 0.35$ $h^{-1}$, this condition is readily satisfied.

***Effective Growth Rate.***

When cell densities are spatially varying, even if metabolite concentrations are initially uniform, they will acquire spatial structure over time. Therefore, the growth rate for each strain, which is a function of the local concentrations of metabolites at each time, will vary spatially and temporally. To assess the overall or effective growth rate of each strain at each time within the experimental system, we introduce a spatially averaged growth rate, weighted by the cell densities. For the LOF mutant and ancestor, they are given by

$$\mu^b_{\text{effective}}(t) = \frac{\int_{\mathcal{V}} \mu^b(G(\vec{r},t),A(\vec{r},t))\, c_b(\vec{r},t)\, d^3r}{\int_{\mathcal{V}} c_b(\vec{r},t)\, d^3r} \quad , \tag{22}$$

and

$$\mu^a_{\text{effective}}(t) = \frac{\int_{\mathcal{V}} \mu^a(G(\vec{r},t),A(\vec{r},t),N(\vec{r},t))\, c_a(\vec{r},t)\, d^3r}{\int_{\mathcal{V}} c_a(\vec{r},t)\, d^3r}, \tag{23}$$

where the integration is over the total spatial domain $\mathcal{V}$ (in our case, the closed experimental chamber); $\mu^b\big(G(\vec{r},t),A(\vec{r},t)\big)$ is the LOF mutant growth rate, and $\mu^a\big(G(\vec{r},t),A(\vec{r},t),N(\vec{r},t)\big)$ given by

$$\mu^a\big(G(\vec{r},t),A(\vec{r},t),N(\vec{r},t)\big) = \mu^a_A\big(G(\vec{r},t),A(\vec{r},t)\big) + \mu\big(G(\vec{r},t),A(\vec{r},t),N(\vec{r},t)\big), \tag{24}$$

is the total ancestor growth rate (as the sum of growth rates on $NH_4^+$ and $N_2$), at each point in space at each time.

As metabolites are produced, consumed, and diffuse, the effective growth rates capture the following relevant features of growth and expansion in the coculture (visualized in Fig. S6):

i. Even if the calculated growth rate $\mu^i(\vec{r},t)$ (where $i = a, b$ refers to the ancestor or LOF mutant) is high in a given spatial region but the ancestor/LOF mutant is not present at that location at that time (that is, $c_i(\vec{r},t) \approx 0$), the effective growth rate $\mu^i_{\text{effective}}(t)$ will not be high.
ii. Spatial separation of the LOF mutant and ancestor allows the LOF mutant to grow in regions where it is not directly competing with the ancestor: even if $u^a > u^b$ in such a region, the ancestor does not outcompete the LOF mutant because it is not present to take advantage of this higher growth rate.
iii. Non-motile cells do not diffuse quickly through agarose, avoiding competition, whereas $NH_4^+$ diffuses quickly enough from the producer to support appreciable LOF mutant effective growth in regions where the ancestor is not present.
iv. In the absence of motility, slow diffusion of cells implies that by the time the ancestor reaches spatial pockets where the LOF mutant has experienced growth, glucose may be completely depleted, halting the ancestor's further growth and expansion.

To illustrate these points, in Fig. 6B and C, starting with a small initial frequency for the LOF mutant ($f_0 = 0.061$), we plot cross-sections of the ancestor and LOF mutant cell density profiles ($c_a(\vec{r},t)$ and $c_b(\vec{r},t)$), and growth rates ($\mu^a(\vec{r},t)$ and $\mu^b(\vec{r},t)$), at $y = 1$ cm at $t = 10$ h. We note that on the right side of the chamber where the ancestor is localized, its growth rate is approximately zero given that glucose is depleted. On the left side of the chamber, the ancestor growth rate is high (given the availability of glucose, $NH_4^+$ and $N_2$); however, the ancestor is not present in this region to take advantage of the high growth rate. Therefore, the effective ancestor growth rate at 10 h is small. On the other hand, the peak of the LOF mutant cell density is located in a region where its growth rate is high; consequently, the effective growth rate is comparable and high.

***LOF change in frequency.***

The metric used to quantify emergence of the loss of function mutant in invasion from rare assays is the change in frequency defined as

$$\Delta f(t) = f(t) - f(0), \tag{25}$$

where $f(t)$ is the LOF mutant frequency at time $t$, and $f_0 = f(0)$ is the frequency at $t = 0$. $f(t)$ is given by

$$f(t) = \frac{\mathcal{N}_b(t)}{\mathcal{N}_b(t)+\mathcal{N}_a(t)}, \tag{26}$$

and $\mathcal{N}_{a,b}(t)$ is the total number of ancestor/LOF mutant cells in the chamber at time $t$,

$$\mathcal{N}_{a,b}(t) = \int_{\mathcal{V}} c_{a,b}(\vec{r},t)\ d^3r. \tag{27}$$

Therefore,

$$\Delta f(t) = \frac{\mathcal{N}_b(t)}{\mathcal{N}_b(t)+\mathcal{N}_a(t)} - \frac{\mathcal{N}_b(0)}{\mathcal{N}_b(0)+\mathcal{N}_a(0)}, \tag{28}$$

Note that the time rate of change of total cell numbers is:

$$\frac{d\mathcal{N}_a}{dt} = \int_{\mathcal{V}} \mu^a\big(G(\vec{r},t), A(\vec{r},t), N(\vec{r},t)\big)\ c_a(\vec{r},t)\, d^3r. \tag{29}$$

and

$$\frac{d\mathcal{N}_b}{dt} = \int_{\mathcal{V}} \mu^b\big(G(\vec{r},t), A(\vec{r},t)\big)\ c_b(\vec{r},t)\, d^3\mathrm{r}. \tag{30}$$

From the definition of the effective growth rates, given by Eq. (22)-(23), we have

$$\frac{d\mathcal{N}_{a,b}}{dt} = \mathcal{N}_{a,b}(t)\mu^{a,b}_{\mathrm{effective}}(t), \tag{31}$$

and integrating we can find

$$\mathcal{N}_{a,b}(t) = \mathcal{N}_{a,b}(0)e^{\int_0^t \mu^{a,b}_{\mathrm{effective}}(t')dt'}, \tag{32}$$

where the overall exponential growth of the ancestor/LOF mutant is given by the time integral of the effective growth rate

$$\int_0^t \mu_{\text{effective}}^{a,b}(t')dt'. \tag{33}$$

Therefore, the change in frequency can be written as

$$\Delta f(t) = \frac{\mathcal{N}_b(0)}{\mathcal{N}_b(0)+\mathcal{N}_a(0)\, e^{\int_0^t \left(\mu_{\text{effective}}^a(t')-\mu_{\text{effective}}^b(t')\right)dt'}} - \frac{\mathcal{N}_b(0)}{\mathcal{N}_b(0)+\mathcal{N}_a(0)}. \tag{34}$$

From Eq. (34) we can see that whether the change in frequency, $\Delta f(t)$, is positive or negative depends on the difference in the overall growth. This can be summarized as:

i. Difference in overall growth between ancestor and the LOF mutant is positive:
$$\int_0^t \left(\mu_{\text{effective}}^a(t') - \mu_{\text{effective}}^b(t')\right) dt' > 0 \Rightarrow e^{\int_0^t \left(\mu_{\text{effective}}^a(t')-\mu_{\text{effective}}^b(t')\right)dt'} > 1 \Rightarrow f(t) < f(0),$$

ii. Difference in overall growth between ancestor and the LOF mutant is negative:
$$\int_0^t \left(\mu_{\text{effective}}^a(t') - \mu_{\text{effective}}^b(t')\right) dt' < 0 \Rightarrow e^{\int_0^t \left(\mu_{\text{effective}}^a(t')-\mu_{\text{effective}}^b(t')\right)dt'} < 1 \Rightarrow f(t) > f(0),$$

Figure S7 show the effective growth rates for the LOF mutant and ancestor (top panel for each time point), their time integrals (lower panel for each time point) and the change in frequency (right panel for each time point). The vertical lines in each plot refer to the time point $t$ at which the change in frequency is considered. The shaded area under each curves is the time integral of the effective growth rate up to that time point; this area corresponds to the overall growth at that time, and marked by the dashed line in the lower panels. These figures demonstrate the fact that initially (Fig. S7, 2 h), the overall growth of the ancestor is greater than that of the LOF mutant and the change in frequency is negative; however, at later times (Fig. S7, 12.5 h), as glucose in the vicinity of the localized ancestor cell density becomes depleted (Fig. S6), the overall growth of the LOF mutant surpasses that of the ancestor, and the change in frequency becomes positive. When the overall growths are equal, the change in frequency is zero (Fig. S7, 5.5 h).

**Table S1. Model parameter values.**

| Parameter | Description | Value | Units |
|---|---|---|---|
| Maximum growth rates | | | |
| $\mu_{max,A}^{a}$ | ancestor on $NH_4^+$, measured, this study | 0.348[a] | $h^{-1}$ |
| $\mu_{max,A}^{b}$ | LOF mutant on $NH_4^+$, measured, this study | 0.348 x 1.1, 1.3 | $h^{-1}$ |
| $\mu_{max,A}^{p}$ | producer on $NH_4^+$, [16] | 0.0875 | $h^{-1}$ |
| $\mu_{max,N}^{a}$ | ancestor on $N_2$, measured, this study | 0.222[a] | $h^{-1}$ |
| $\mu_{max,N}^{p}$ | producer on $N_2$, measured [1] | 0.0772 | $h^{-1}$ |
| Metabolite production levels and rates | | | |
| $F_A^{p,a}$ | $NH_4^+$ excretion level by producer, ancestor in privatized model, estimated, this study | $6.5 \times 10^{-10}$ | µmol $cell^{-1}$ |
| $\widetilde{F_A^{p,a}}$ | $NH_4^+$ excretion rate by producer, ancestor in non-privatized model, estimated, this study | $6.5 \times 10^{-10}$ | µmol $cell^{-1}$ $s^{-1}$ |
| $F_C$ | organic acid production level during growth on $NH_4^+$ by ancestor, LOF mutant, measured, this study | $1.0 \times 10^{-6}$ | µmol $cell^{-1}$ |
| $F_{C,N}^{a}$ | organic acid production level during growth on $N_2$ by ancestor, measured, this study | $1.95 \times 10^{-6}$ | µmol $cell^{-1}$ |
| Half-saturation constants | | | |
| $K_A^{a,b}$ | ancestor, LOF mutant affinity for $NH_4^+$, assumed [17] | 0.01 | mM |
| $K_A^{p}$ | producer affinity for $NH_4^+$, assumed [18] | 0.1 | mM |
| $K_G^{a,b}$ | ancestor, LOF mutant affinity for glucose, assumed [19] | 0.02 | mM |
| $K_C^{p}$ | producer affinity for organic acids, assumed | 0.01 | mM |
| $K_N^{a,p}$ | ancestor, producer affinity for $N_2$, measured [20] | 6 | mM |
| Growth yields | | | |
| $Y_N^{p}$ | producer from $N_2$, measured [20] | $5 \times 10^{8}$ | cell $µmol^{-1}$ |
| $Y_N^{a}$ | ancestor from $N_2$, measured | $4 \times 10^{8}$ | cell $µmol^{-1}$ |
| $Y_G$ | ancestor, LOF mutant from glucose with $NH_4^+$, measured | $1.9 \times 10^{6}$ | cell $µmol^{-1}$ |
| $Y_{G,N}^{a}$ | ancestor from glucose with $N_2$, measured | $1 \times 10^{6}$ | cell $µmol^{-1}$ |
| $Y_A^{a,b}$ | ancestor, LOF mutant from $NH_4^+$, measured | $2 \times 10^{8}$ | cell $µmol^{-1}$ |
| $Y_A^{p}$ | producer from $NH_4^+$, assumed | $5 \times 10^{8}$ | cell $µmol^{-1}$ |
| $Y_{C,N}^{p}$ | producer from organic acids with $N_2$, measured | $1.3 \times 10^{8}$ | cell $µmol^{-1}$ |
| $Y_{C,A}^{p}$ | producer from organic acids with $NH_4^+$, assumed | $1.3 \times 10^{8}$ | cell $µmol^{-1}$ |
| Diffusion constants in liquid | | | |
| $D_a, D_b, D_p$ | non-motile ancestor, LOF mutant, producer cells | $1.8 \times 10^{-5}$ | $cm^2$ $h^{-1}$ |
| $D_C, D_G, D_A, D_N$ | organic acids, glucose, $NH_4^+$, $N_2$ | 0.03 | $cm^2$ $h^{-1}$ |
| Diffusion constants in 0.15% agarose | | | |
| $D_a, D_b, D_p$ | non-motile ancestor, LOF mutant, producer cells | $6 \times 10^{-6}$ | $cm^2$ $h^{-1}$ |
| $D_C, D_G, D_A, D_N$ | organic acids, glucose, $NH_4^+$, $N_2$ | 0.01 | $cm^2$ $h^{-1}$ |
| Default initial values | | | |
| $C$ | organic acids | 0.000001 | mM |
| $G$ | glucose | 5 | mM |
| $A$ | $NH_4^+$ | 0.00005 | mM |
| $N$ | $N_2$; assumed to be fully dissolved | 70 | mM |
| $c_i$ | cell density | varied[b] | cells $ml^{-1}$ |

[a] To avoid our interpretations being influenced by the extreme growth rates of *V. natriegens* we ran simulations with values corresponding to a slower-growing fermentative $N_2$-fixing bacterium, *Zymonomas mobilis* [21].

[b] Simulated initial cell densities were the same as those measured in experimental assays.

**Table S2. Strains**

| Strain | Designation; genotype; phenotype | Source or reference |
|---|---|---|
| *R. palustris* strains | | |
| CGA0092 | Wild-type; spontaneous $Cm^R$ derivative of CGA001 | [22, 23] |
| CGA4039 | Δnitrogenase or $ΔN_2$ase; nifA*, Δ*uppE* (RPA2750), Δ*nifH*, Δ*vnfH,* Δ*anfH*; incapable of $N_2$ fixation | This study |
| CGA4005 | *nifA**, Δ*hupS*, Δ*uppE*; constitutive nitrogenase expression resulting in $NH_4^+$ excretion, incapable of $H_2$ oxidation, decreased capacity for biofilm formation | [1] |
| CGA4067 | Producer; *nifA**, Δ*hupS*, Δ*uppE*, Δ*motAB*; $NH_4^+$ excretion, incapable of $H_2$ oxidation, decreased capacity for biofilm formation, non-motile due to absence of flagellar stator | This study |
| *V. natriegens* strains | | |
| TND1964 | Wild type (WT); ATCC14048, pMMB-tfoX | [24] |
| NH003 | WT Δ*dns*::$Kan^R$, pMMB-tfoX | This study |
| NH004 | WT Δ*dns*::$Spec^R$, pMMB-tfoX | This study |
| OFS003 | Ancestor; Δ*dns*::$Kan^R$, pMMB-tfoX, Δ*flgE,*; non-motile due to the absence of the flagellar hook | This study |
| OFS004 | LOF mutant; Δ*dns*::$Spec^R$, pMMB-tfoX, Δ*flgE,* Δ*nifA*; non-motile and incapable of $N_2$ fixation | This study |
| OFS005 | pMMB-tfoX, Δ*nifA::*$Spec^R$; incapable of $N_2$ fixation | This study |

**Table S3. Plasmids**

| Plasmid | Description | Source or reference |
|---|---|---|
| pJQ200SK | Suicide vector used for making deletion mutants in *R. palustris* | [25] |
| pJQΔ*motAB* | Suicide vector for deleting the flagellar stator genes in *R. palustris* | [26] |
| pJQΔ*nifH* | Suicide vector for deleting the Mo-nitrogenase gene in *R. palustris* | [27] |
| pJQΔ*vnfH* | Suicide vector for deleting the V-nitrogenase gene in *R. palustris* | [27] |
| pJQΔ*anfH* | Suicide vector for deleting the Fe-nitrogenase gene in *R. palustris* | [27] |
| pMMB-tfoX | Expression vector with IPTG-inducible *V. cholerae tfoX* to allow for natural transformation. | [24] |

**Table S4. Primers**

| Primer | Sequence | Description | Source or reference |
|---|---|---|---|
| OFS64 | gatatcgaaatgccagagatggac | Δ*flgE* forward, 3 kb upstream | This study |
| OFS65 | gtcgacggatccccggaattccaaagtct cctgatctcgc | Δ*flgE* MUGENT MiniFRT overlap reverse | This study |
| OFS66 | agaagcagctccagcctacaatcgcga gccggataatc | Δ*flgE* MUGENT MiniFRT overlap forward | This study |
| OFS67 | acggttgggttattaccgacaatc | Δ*flgE* reverse, 3 kb downstream | This study |
| OFS70 | gacttgcagaagtattggaagtcg | Δ*flgE* sequencing forward | This study |
| OFS71 | gtcataaagtaatcaagtgagctgc | Δ*flgE* sequencing reverse | This study |
| JPL007 | gagataagtgacgcaagccg | Δ*nifA* reverse, 3 kb downstream | This study |
| JPL006 | gaagcagctccagcctacacaaggctg ctcaatatgacgc | Δ*nifA* MUGENT MiniFRT overlap forward | This study |
| JPL005 | gtcgacggatccccggaatccgccaata actgacgctc | Δ*nifA* MUGENT MiniFRT overlap reverse | This study |
| SB001 | ggacatgattcactcctaaaccg | Δ*nifA* forward, 3 kb downstream | This study |
| JPL48 | atgatagaagatcattctc | Δ*nifA* sequencing forward | This study |
| JPL49 | tcagatttgtttcatctcgatatt | Δ*nifA* sequencing reverse | This study |
| NH022 | cgaggtgaagatcattcatttcc | Δ*dns* forward, 1 kb upstream | This study |
| NH025 | cttagtgattgggtcactcattgg | Δ*dns* reverse, 1 kb upstream | This study |
| NH014 | ctctgcaccactaccgtc | Δ*dns* sequencing forward | This study |
| NH015 | cgaataccgatgtcgctgc | Δ*dns* sequencing reverse | This study |
| Prm361 | attccggggatccgtcgacctgcagttca gaagcagctccagcctaca | miniFRT cassette forward | [28] |
| Prm362 | tgtaggctggagctgcttctgaactgcag gtcgacggatccccggaat | miniFRT cassette reverse | [28] |

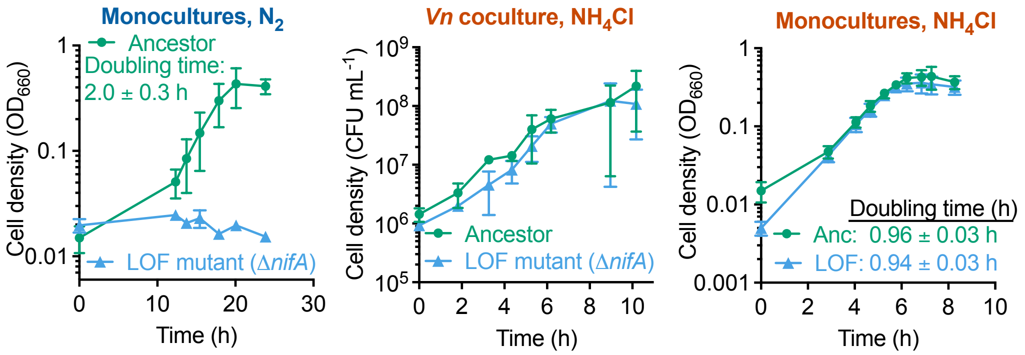


**Figure S1.** The *V. natriegens* Δ*nifA* LOF mutant cannot use $N_2$ but the ancestor (Anc) and LOF mutant growth rates are similar with $NH_4Cl$; (A) monoculture growth with $N_2$ as the nitrogen source; data points and doubling time are means ± SD, n = 3; (B) growth of the ancestor and the LOF mutant in cocultures with 10 mM $NH_4Cl$; data points are means ± SD n = 3; (C) monoculture growth with 10 mM $NH_4Cl$; data points and doubling time are means ± SD, n = 3; p = 0.59 (two-tailed t-test).

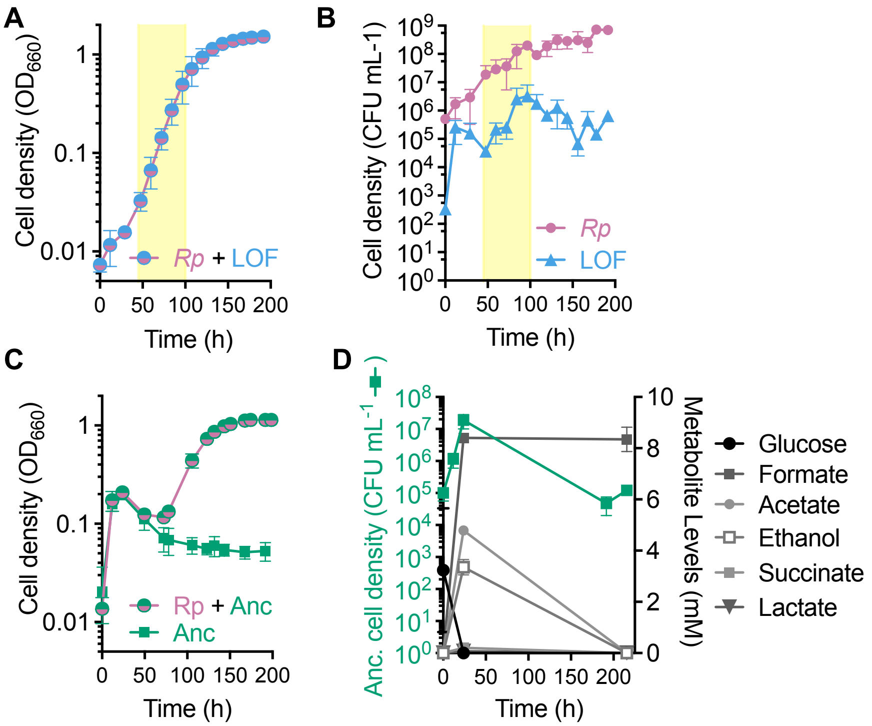


**Figure S2.** *R. palustris* Nx 'producer' (*Rp*) and the *V. natriegens* Δ*nifA* LOF mutant grow by reciprocal cross-feeding; (A, B) growth of cocultures pairing the producer and the LOF mutant, tracked using coculture turbidity (A) or colony forming units for each population (B); the exponential phase is highlighted in yellow; (C) comparison of growth curves between WT *V. natriegens* 'ancestor' (Anc) monocultures and cocultures pairing the ancestor with the producer; (D) glucose and fermentation product concentrations in ancestor + producer cocultures; (A-D) data points are means ± SD, n = 3.

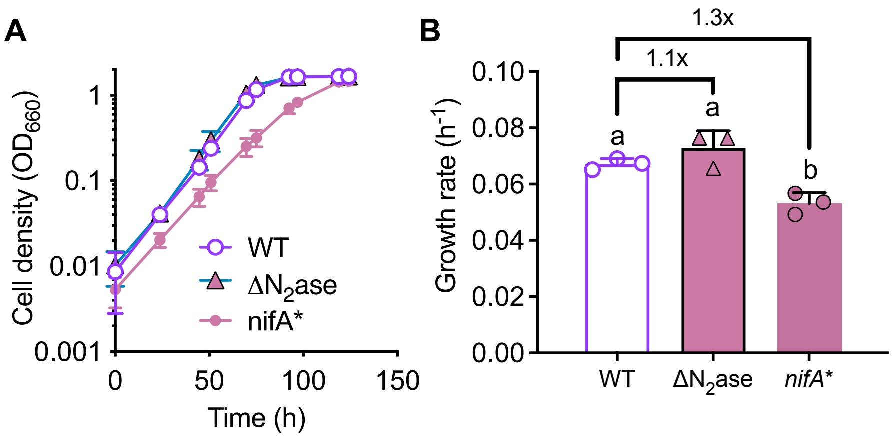


**Figure S3.** Estimation of growth advantage imparted by loss of nitrogenase (A) growth of *R. palustris* CGA0092 (WT), CGA4039 (Δnitrogenase; Δ$N_2$ase) (CGA4005 (*nifA**) in media with $NH_4Cl$; (B) growth rates calculated from the exponential phase in (A); the floating values indicate fold-differences in growth rate; floating letters indicate significant differences according to a one-way ANOVA with Tukey's multiple comparisons test ($p >$ 0.05); a similar comparison with *V. natriegens* was not possible as we have yet to successfully generate a NifA* strain.

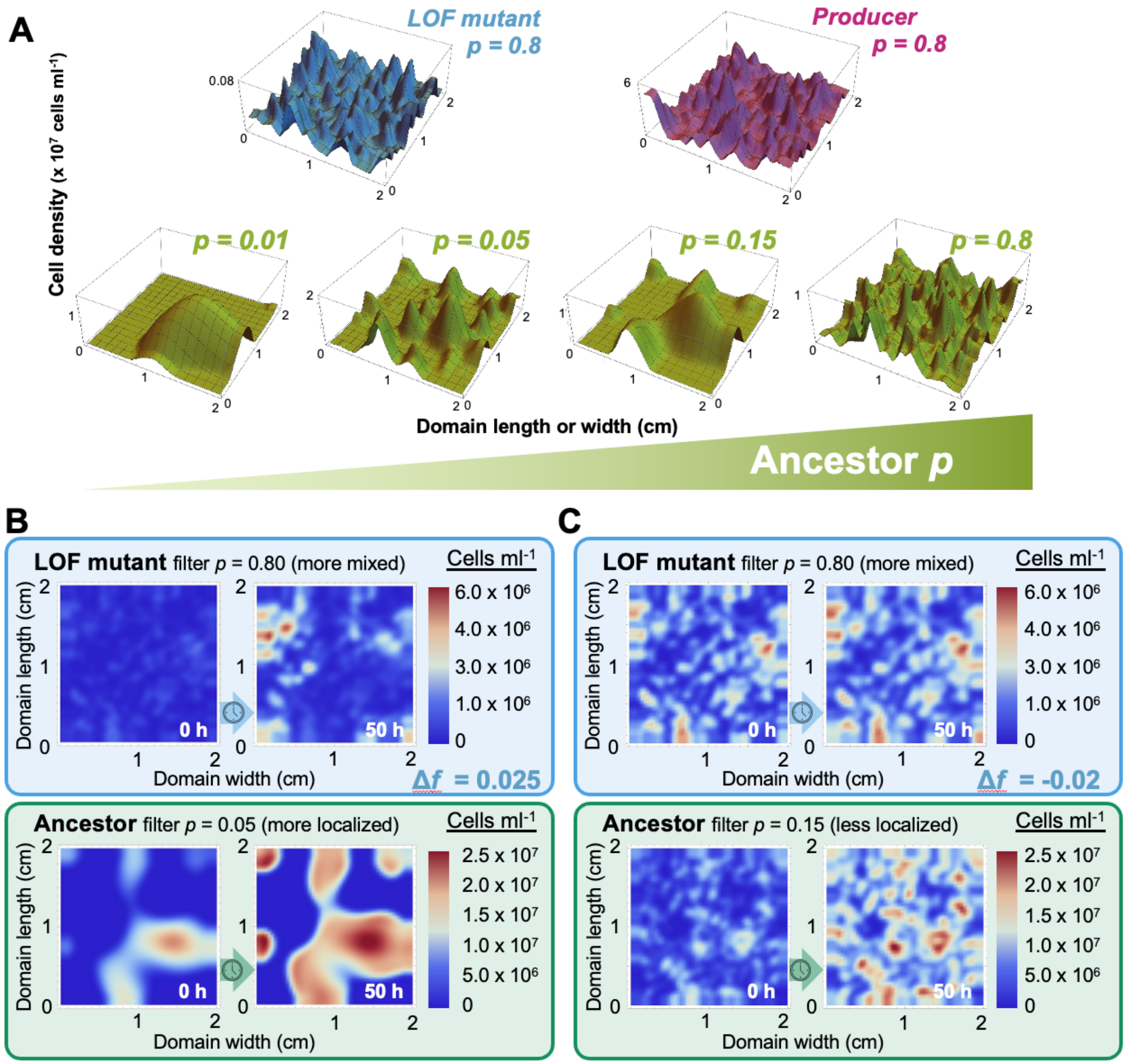


**Figure S4.** The degree to which initial ancestor spatial cell densities are randomized affects whether the LOF mutant is enriched; (A-C) initial LOF mutant frequency $f_0 = 0.061$; LOF mutant and producer initial cell densities were held fixed with a filter parameter $p = 0.8$; (A) upper row, an example of randomized initial LOF mutant and producer cell densities; lower row, an example of how the same ancestor initial cell density is randomly distributed using different spatial filter values of $p$; the ancestor spatial cell density becoming more fine grained and more mixed with the other population with increasing $p$; (B-C) examples of how a low ancestor spatial filter (B, $p = 0.05$, more localized) can lead to LOF mutant enrichment whereas a higher ancestor spatial filter (C, $p = 0.15$, less localized), prevents LOF mutant enrichment; panel B is the same data as in Fig. 4D.

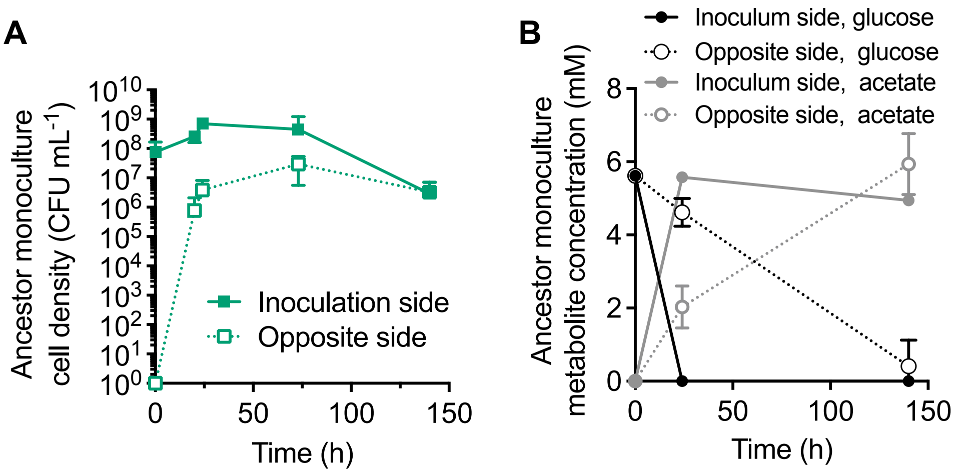


**Figure S5.** Population and metabolic trends are slower on the opposite side of the vial from the inoculation site in *V. natriegens* ancestor monocultures in 0.15% agarose media. (A) ancestor growth inferred from samples taken from the inoculation site and opposite side of the vial; (B) metabolite levels inferred from samples taken from the inoculation site and opposite side of the vial; (A, B) points are means ± SD, n = 3.

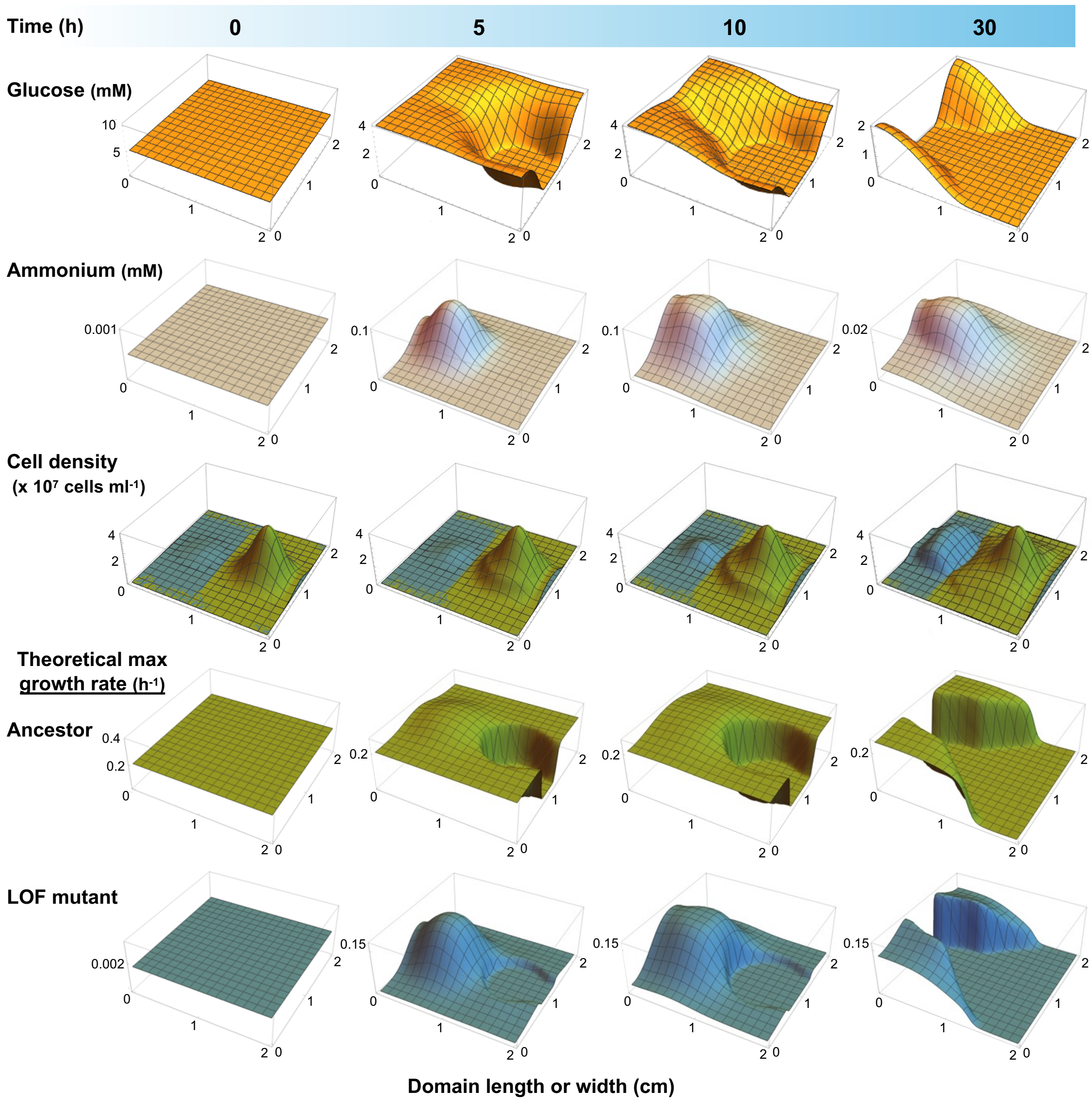


**Figure S6.** Snapshots from simulations where the producer and LOF mutant (blue) are colocalized away from the ancestor (green) (σ = 0.2). Glucose is quickly depleted at the ancestor inoculation site (5 h). The LOF mutant consumes enough glucose near its inoculation site to slow the advance of the ancestor into its territory (10 h). Both populations the expand to the sides where glucose remains. Note that theoretical maximum growth rates only show what is possible at a given location (regions with few if any cells can coincide with a high theoretical maximum growth rate.

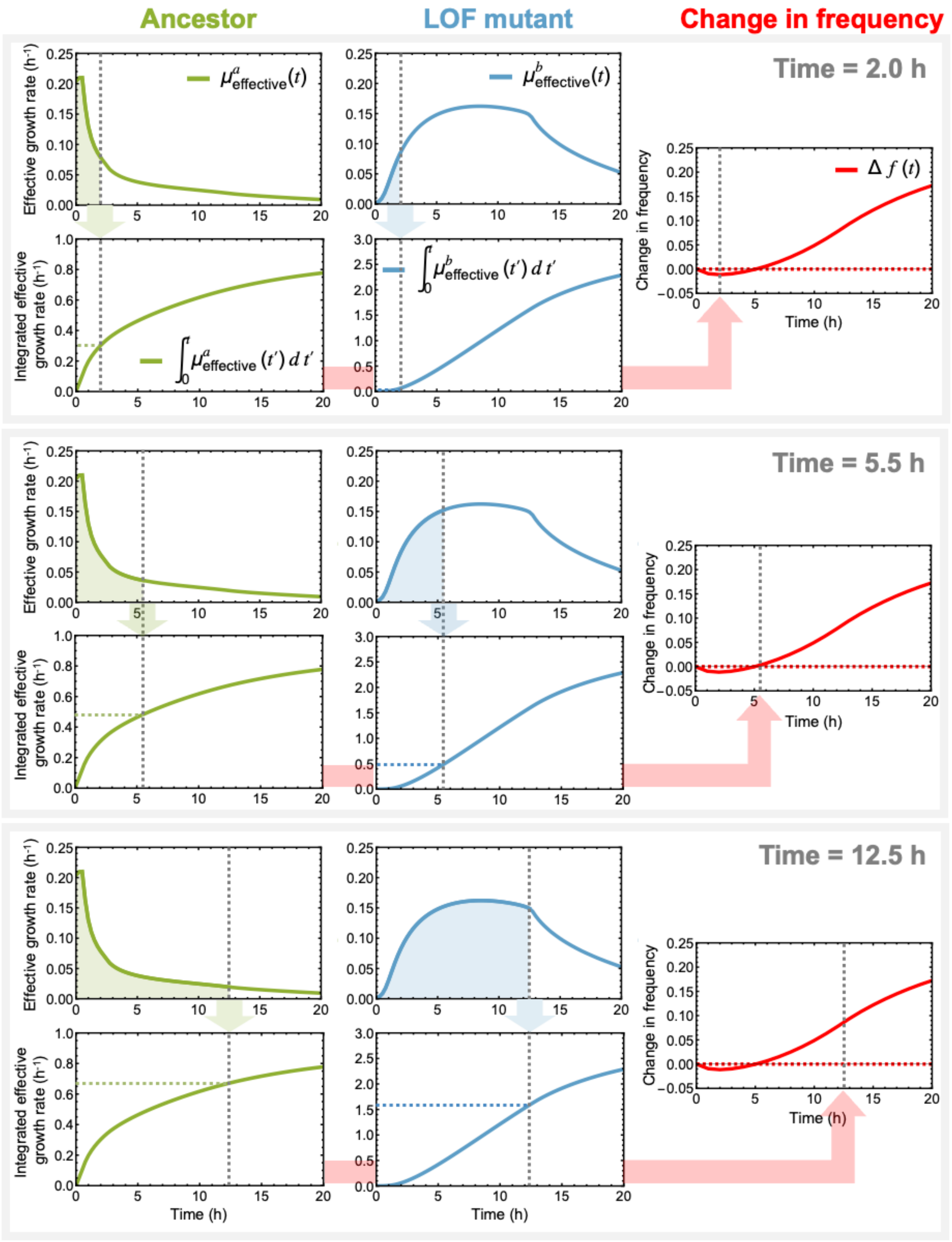


**Figure S7.** Change in LOF mutant frequency ($\Delta f(t)$) can be calculated by integrating the effective growth rate of each *V. natriegens* population; three time points are shown (2.0, 5.5, and 12.5 h) to illustrate how the change in frequency plot is built; shaded regions are the integrated effective growth rates up to that time point, leading to a data point in the graph below (follow the shaded green and blue arrows); the integrated effective growth rates are then used to calculate the change in frequency (follow the shaded red arrows); for more details see the description of equations 33 and 34, above.

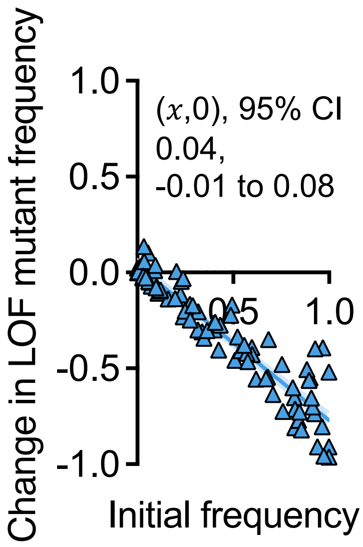


**Figure S8.** Compilation of all data from every IFR experiment performed in static liquid; each point is a single experimental coculture; initial frequency range = 0.1% - 99.9% using *V. natriegens* populations only; $x$-intercept and 95% CI error bands were determined using linear regression analysis.

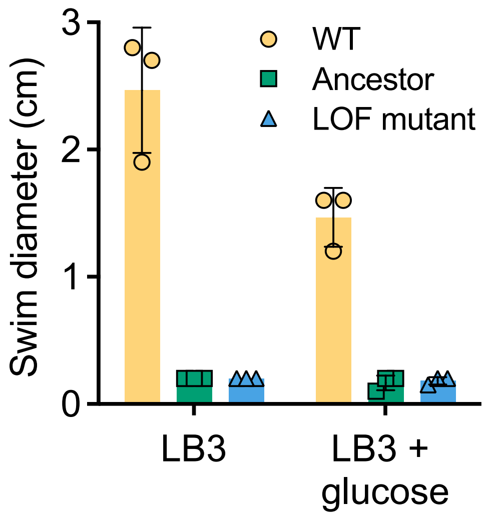


**Figure S9.** Deleting *V. natriegens flgE* eliminates motility; two kinds of media were used to assess whether 10 mM glucose would fully inhibit motility; bars are the mean of three biological replicates ± SD. WT, NH003.

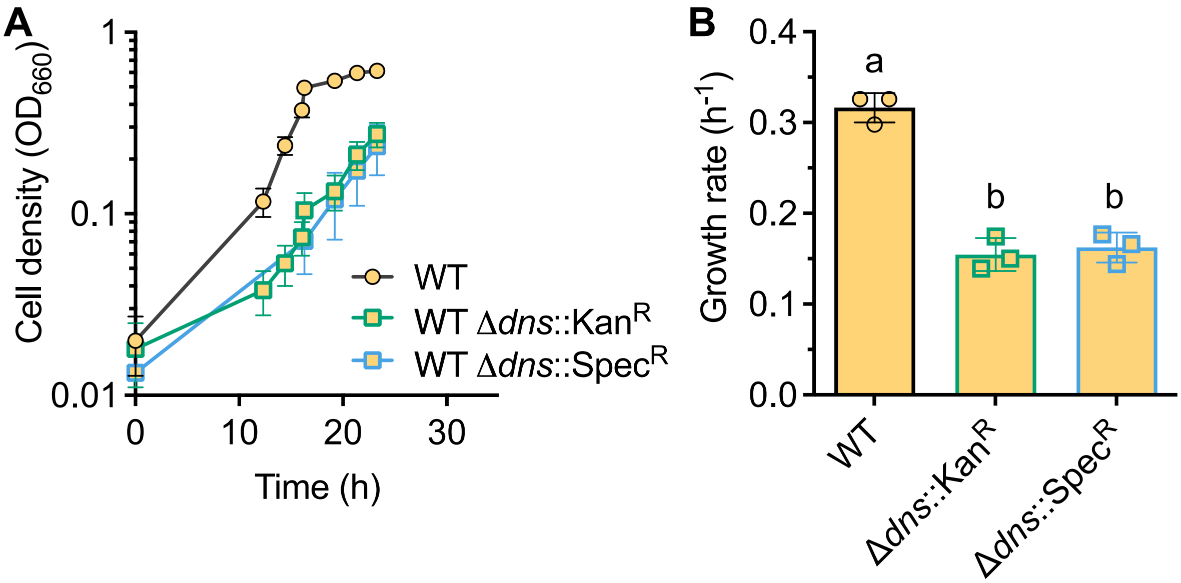


**Figure S10.** Δ*dns*::Kan$^R$ and Δ*dns*::Spec$^R$ have equal effects on growth rate; (A) *V. natriegens* monoculture growth curves when *dns* is intact or replaced with different antibiotic cassettes; (B) monoculture growth rates from (A); floating letters indicate significant differences from a one-way ANOVA with Tukey's multiple comparison post-test ($p > 0.05$); (A, B) points or bars are means ± SD, n = 3.

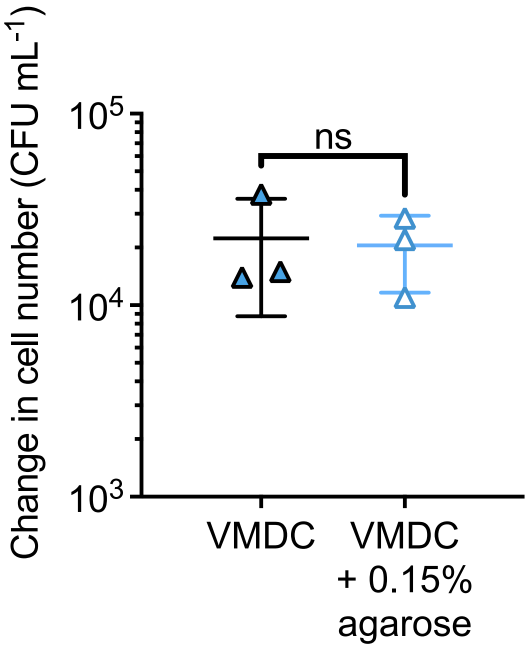


**Figure S11.** Washed 0.15% agarose does not contain enough nitrogen to support growth of the *V. natriegens* LOF mutant (OFS005); points are individual cultures; bar is mean ± SD, n=3; ns = non-significat based on a two-tailed unpaired t-test ($p$ = 0.85).
